# Nuclear envelope budding is a response to cellular stress

**DOI:** 10.1101/413153

**Authors:** Dimitra Panagaki, Jacob T. Croft, Katharina Keuenhof, Lisa Larsson-Berglund, Stefanie Andersson, Verena Kohler, Sabrina Büttner, Markus J. Tamás, Thomas Nyström, Richard Neutze, Johanna L. Höög

**Author notes:** Johanna L. Höög, **Email:**. These authors contributed equally to the manuscript (Dimitra Panagaki, Jacob T. Croft).

## Abstract

Nuclear envelope budding (NEB) is a recently discovered alternative pathway for nucleocytoplasmic communication distinct from the movement of material through the nuclear pore complex. Through quantitative electron microscopy and tomography, we demonstrate how NEB is evolutionarily conserved from early protists to human cells. In the yeast *Saccharomyces cerevisiae*, NEB events occur with higher frequency during heat shock, upon exposure to arsenite or hydrogen peroxide, and when the proteasome is inhibited. Yeast cells treated with azetidine-2-carboxylic acid, a proline analogue that induces protein misfolding, display the most dramatic increase in NEB, suggesting a causal link to protein quality control. This link was further supported by both localization of ubiquitin and Hsp104 to protein aggregates and NEB events, and the evolution of these structures during heat shock. We hypothesize that NEB is part of normal cellular physiology in a vast range of species and that in *S. cerevisiae* NEB comprises a stress response aiding the transport of protein aggregates across the nuclear envelope.

**Significance Statement:** A defining feature of eukaryotes is the nuclear envelope, a double lipid bilayer that serves to isolate and protect the cells genetic material. Transport of large molecules over this barrier is believed to occur almost exclusively *via* the nuclear pores. However, herpes virions and mega ribonucleoproteins (megaRNPs) use an alternative means of transport – *via* nuclear envelope budding (NEB). Here, we show NEB is a ubiquitous eukaryotic phenomenon and increases when exposed to various forms of cellular stress. NEB frequency was maximal when the cell was challenged with a drug that induces protein misfolding, indicating this transport pathway plays a role in protein quality control. These results imply that NEB is an underappreciated yet potentiallyfundamental means of nuclear transport.

## Introduction

The nucleus is the most prominent organelle in eukaryotic cells, enclosing most of the cellular genetic material within a double lipid bilayer called the nuclear envelope. These two membranes arose as a critical evolutionary step distinguishing eukaryotes from prokaryotes and restricting which molecules come into contact with the cellular DNA. As the nuclear envelope is not permeable to most of the molecules inside the cell, special structures called nuclear pore complexes (NPCs) exist on its surface that allow highly selective translocation over the nuclear membrane (1–3). Molecules smaller than 30-40 kDa can passively penetrate the NPC from the cytoplasm into the nucleus and *vice versa*, whereas bigger molecules require interaction with nuclear transport receptors and form importin/exportin complexes that are guided through the pores (4–8).

Since its discovery in 1954, the NPC has generally been accepted as the only means of communication between the cytoplasm and the nucleoplasm (9). However, herpes simplex virus replicates in the nucleoplasm and is released into the cytosol via an outwards budding of the nuclear envelope (10–13). This demonstrates another pathway for nuclear export and there have been several observations suggesting that nuclear envelope budding (NEB) also occurs in healthy cells, with different interpretations of this mechanism being suggested (14–27). These observations were made in a diverse set of organisms but primarily using cells that are under differentiation (embryonic cells) and various terminologies were given to describe the NEB process (Supplementary Table 1). The most recent reports were made in healthy developing *Drosophila melanogaster* larvae, in the sea urchin gastrula and in the budding yeast *Saccharomyces cerevisiae* (28–32). Previous studies have shown that budding events in *D. melanogaster* contain large ribonucleoprotein granules but whether this function is consistent in other cell types remains unknown (28, 31).

Evidence that material can be exported from the nucleus through NEB has sparked speculation that other large cargoes, such as protein aggregates, could be removed from the nucleus by a similar mechanism (33). Protein misfolding can be highly toxic to the cell and therefore multiple stress-induced mechanisms have evolved to cope with proteotoxicity (34–36). Cells experiencing large-scale protein misfolding exhibit DNA mutagenesis, which is one of the first steps towards carcinogenesis (37), demonstrating the importance of protein quality control in the nucleus (38). Despite the cell’s primary protein quality control factors, such as chaperones and the ubiquitin-proteasome system, being active in the nucleus, protein aggregates still form under stressful conditions such as heat shock, because these processes cannot fully cope with the quantity of misfolded proteins (39). In most occasions, protein aggregates will enter the nucleus for degradation but cases of nuclear misfolded proteins being transported out of the nucleus to be degraded in the cytoplasm have also been reported (39). Nuclear protein aggregates greatly exceed the 39 nm size limit of active transportation through the NPCs (40, 41), implying that an alternative pathway for their export is required if they are to be transported over the nuclear envelope.

As well as transporting material from the nucleus to the cytoplasm, NEB can also result in the transfer of a portion of the inner nuclear membrane (INM) to the outer nuclear membrane (ONM), which is continuous with the endoplasmic reticulum (ER). In addition to protein quality control mechanisms in the nucleus and cytoplasm, the endoplasmic reticulum (ER) is equipped with its own set of degradation pathways, to contend with the large quantity of newly synthesized proteins entering the ER (42). Protein degradation in the ER is highly dependent on the ubiquitin proteasome system and facilitates removal of both misfolded soluble and membrane proteins (43). Similarly, degradation of INM proteins can occur via biochemically similar pathways to ER-associated degradation of membrane proteins (44). Several branches of INM-associated degradation (INMAD) exist, each relying on a different E3 ubiquitin ligase to recognize and tag misfolded substrates through ubiquitination steps(45–47). These reflections pose the question: does NEB provide an escape valve needed for clearance of aggregated nuclear proteins? To address this issue, we have examined the NEB pathway of *S. cerevisiae* under five different stress conditions, all of which caused an increase in NEB events. We also show the presence of NEB in five different evolutionarily distant organisms (*Homo sapiens, Caenorhabditis elegans, Saccharomyces cerevisiae, Schizosaccharomyces pombe, Trypanosoma brucei*) under normal growth conditions, which reveals the evolutionary conservation of the NEB pathway. Our findings provide evidence that NEB is part of the cell’s natural stress response and an important pathway to consider when studying nuclear transport, especially with regard to the protein quality control system.

## Results

### NEB increases in frequency during heat shock in *S. cerevisiae*

*S. cerevisiae* provides a well-understood model system, perfectly tailored to investigate if NEB is an active part of the cellular stress response. An easily applied stress impulse is heat-shock and we therefore subjected *S. cerevisiae* cells to a mild constant heat stress (38°C) for a time course of up to 90 minutes (Figure 1**a**). Cells were then cryo-immobilized for electron microscopy studies, which are able to simultaneously visualize both NEB events and protein aggregates, the latter appearing as electron dense content (EDC) (48) within the nucleus (Figure 1**b**). For each time point between 60 to 80 electron micrographs were acquired, each from a randomly chosen cell section that passed through the cell nucleus. NEB events were identified as electron dense material localized between the two nuclear envelopes, causing it to deform, with a small number of events showing budding of the nuclear membrane but seeming to lack an electron dense cargo (S1). A membrane bilayer was often, but not always, detectable around this material. NEB events commonly had an internal texture resembling the nucleoplasm (Figure 1**b**),but in some rare events the bud appeared to contain cytoplasmic material including ribosomes (Figure S1). In one case, a vacuole was seen engulfing a protrusion of the nucleus. This phenomenon is termed piecemeal microautophagy of the nucleus and has previously been described in nutrient deprived cells (49). Piecemeal microautophagy has a distinct morphology from the NEB events and were not included in our quantification.

**Figure 1.**
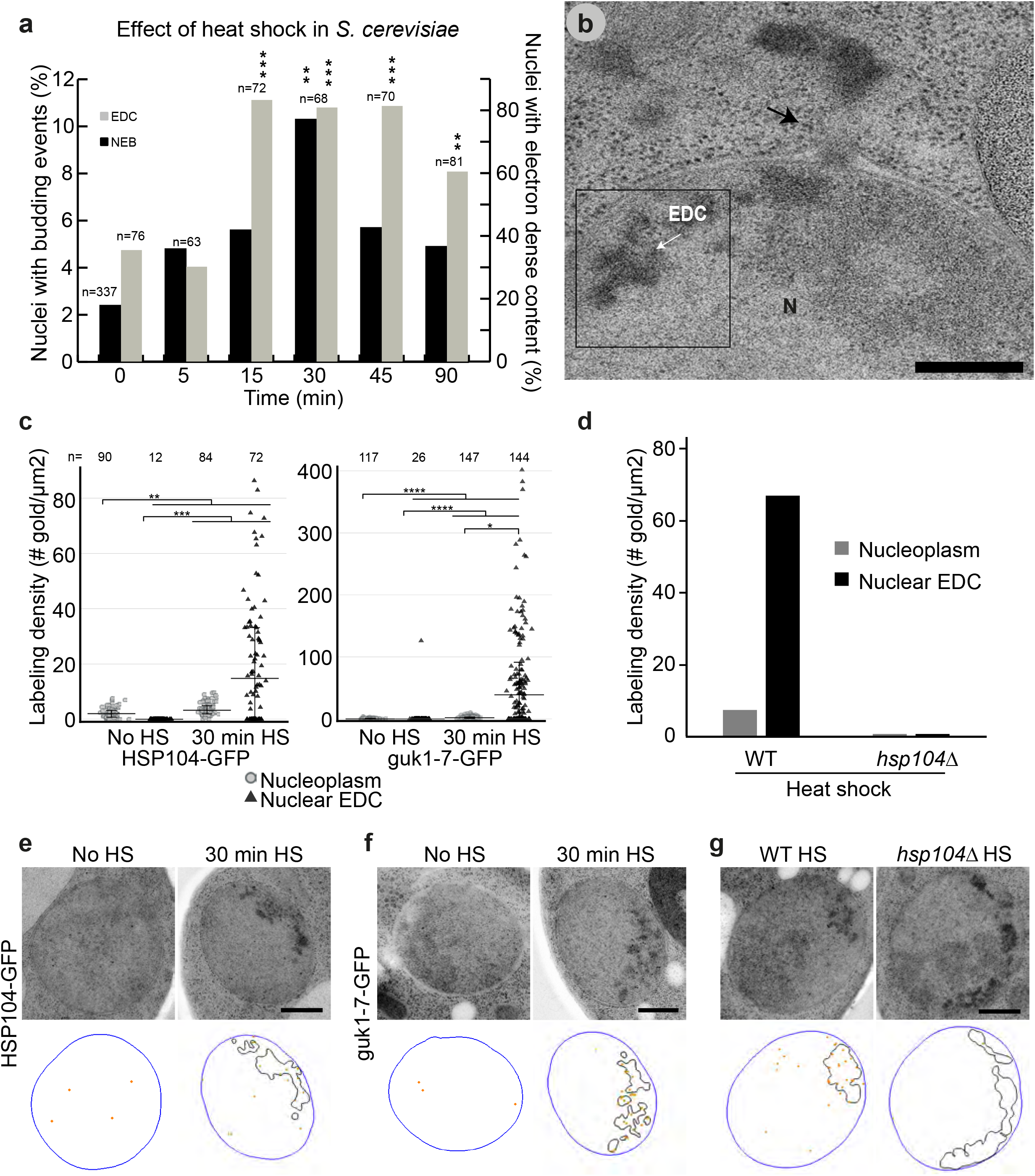
NEB increases in frequency during heat shock in *S. cerevisiae*. (**a**) Percentage of nuclei exhibiting NEB (left axis, black bars) and EDC (right axis, grey bars) as a function of time after cells have been subjected to heat shock (38 °C, n=63-337). NEB peaked at a level of 10.3% after 30 minutes and EDC exhibited a stable plateau of about 81% between 15 and 45 minutes. (**b**) Thin section containing NEB and EDC in heat shocked *S. cerevisiae* cells. Black arrow indicates NEB event, where the outer nuclear membrane clearly protrudes outwards and a particle resides within the perinuclear space. (**c**) Labeling density of gold particles between untreated and heat shocked cells (38 °C for 30 minutes) using an anti-GFP antibody. Both Hsp104-GFP and guk1-7-GFP were preferentially localized at the EDC areas compared to the rest of the nucleoplasm. (**d**) Labeling density of Hsp104 using an anti-Hsp104 antibody in wild type and *hsp104Δ* cells subjected to heat shock at 38 °C for 45 minutes. No gold particles were detected in the mutant strain showing the specificity of the antibodies. (**e**-**g**) Representative images and respective models of the immuno-stained samples quantified in c and d. Hsp104 and guk1-7 are significantly localized to the EDC area whereas only one gold is detected in the EDC of the *hsp104Δ* mutant. Scale bars: 300 nm (**b**) and 500 nm (e, f, g). Abbreviations: NEB, nuclear envelope budding; EDC, electron dense content; N, nucleus; V, vacuole;. \**P*<.05, \*\**P*<.01, \*\*\**P*<.001 vs. zero time point or control group; ns no significant differences between groups.

Morphological variations of NEB (Figure S1) could represent different stages of the same process or different processes. Images were scored both for the presence of NEB events and EDC in the nucleus. Both NEB and EDC increased during heat shock, prior to decreasing near the end of the time course, which potentially reflects a cellular adaptation to the new temperature (Figure 1**a**). EDC reaches a maximum (83.3% of sections) after 15 minutes and stays relatively stable at this level until 90 minutes, at which point it is only present in 60.5% of sections (Figure 1**a**). NEB events were found in 2.4% of cellular sections in undisturbed cultures but this fraction significantly increased to 10.3% after 30 minutes of heat shock. The effect of a stronger heat shock treatment (42°C for 30 minutes) was examined to detect a possible correlation of the NEB frequency to the level of stress. NEB frequency was also here significantly increased in comparison to the control as 16.6% of the nuclear sections displayed NEB (Figure S2).

To verify that the observed EDC in the nucleus are protein aggregates, immuno-electron microscopy against the disaggregase Hsp104 (50) and the model misfolding protein temperature sensitive mutant guk1-7 (51) were performed. In these experiments, the secondary antibody is coupled to a gold particle for visualization using electron microscopy. Both Hsp104 and guk1-7 localised to the EDC material upon heat shock (Figure 1**c**, **e-f**, n = number of examined structures). As a control, an anti-Hsp104 antibody was used to label the EDC in heat shocked wild type and *hsp104Δ* cells to validate the specificity of the labeling. High labeling density of the EDC could be detected in the wild type strain, whereas in the *hsp104Δ* strain almost no unspecific gold particles were found (Figure **d,g**).

The concurrent increase in frequency of NEB and protein aggregation strengthens the hypothesis that NEB may indeed have an important function during normal cellular stress response. This apparent correlation between the frequency of NEB with heat shock motivated us to test if this pathway is solely affected by heat-shock or if cells react in a similar fashion under different stress conditions.

### Hydrogen peroxide and sodium arsenite stressors also increase NEB frequency

Cells may encounter a variety of stress conditions and each of them has a different impact on the cell (52). To investigate whether additional cellular stresses also correlated with an increased frequency of NEB events we challenged *S. cerevisiae* cells with multiple stress stimuli. A natural stimulus is aging which causes a significant induction of stress response pathways in *S. cerevisiae* (35, 53), and has previously been seen to trigger an increase in “nuclear herniations” (54). Several factors are responsible for a measurable decline in cellular and physical functions of aged cells, with one of them being aggregated misfolded proteins (55). We isolated replicatively aged biotinylated *S. cerevisiae* cells, using Streptavidin magnetic beads, from the young unbound population (56) and cryo-immobilized the cells. In old yeast cells, the frequency of NEB events was found to be 9% (Figure 2**a**, 2**d**; n=200 sections) which is significantly higher than an undisturbed *S. cerevisiae* culture (2.4%; n=337) (Figure 1**a**). However, the remaining younger population also displayed a significant increase in NEB events to 6.9% (n=204 sections; Figure 2**d**), indicating that these cells may also have been stressed by the mechanical handling steps associated with isolating the old cells. Thus, we could not confirm the previous finding of significantly increased NEB events in old cells in comparison to young cells (54), in this experimental setup.

**Figure 2.**
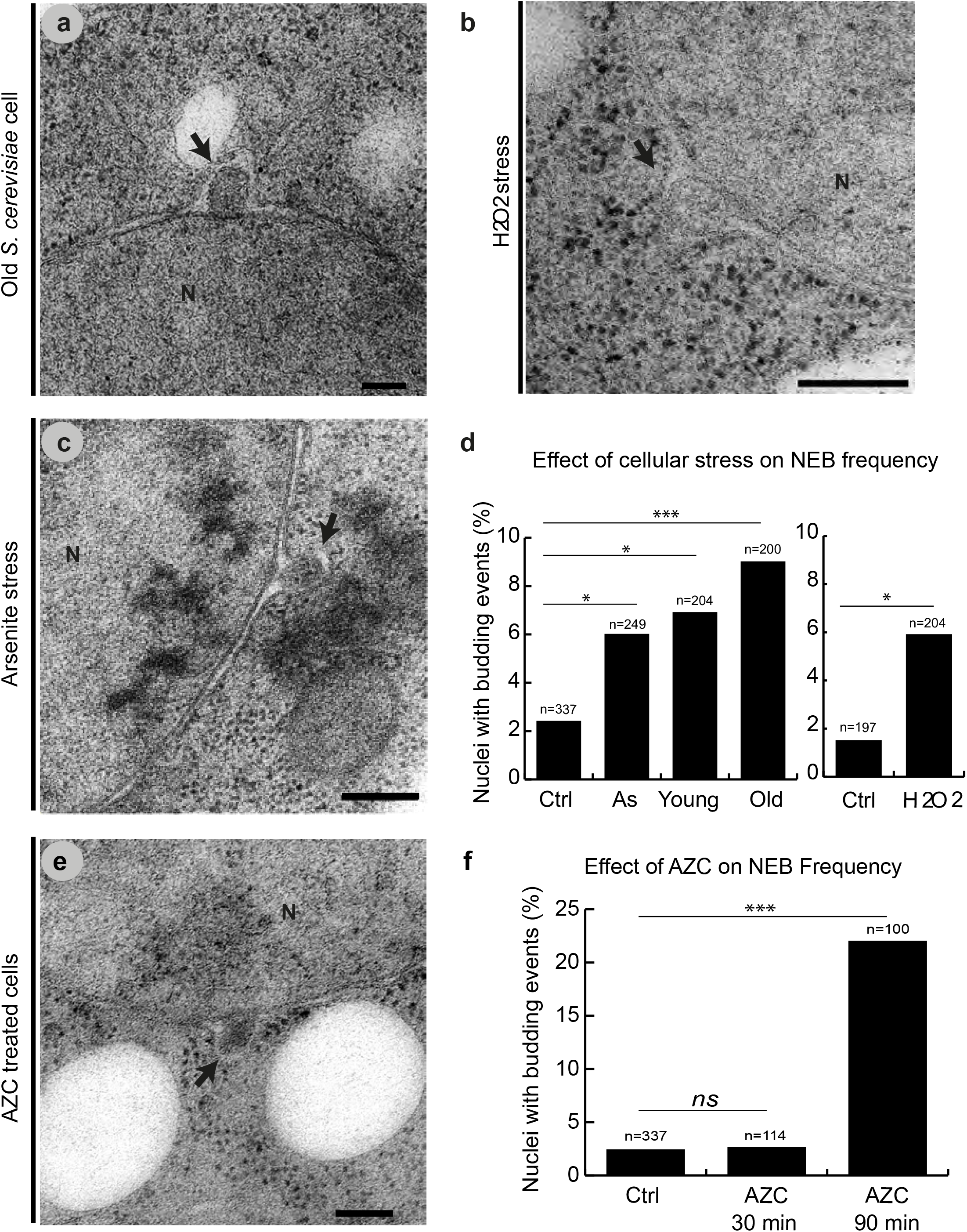
Four different cellular stressors increase NEB frequency in *S. cerevisiae* cells. Thin sections containing NEB events in (**a**) old cells (**b**) cells subjected to oxidative stress with H_2_O_2_ (**c**) cells exposed to sodium arsenite. (**d**) Percentage of nuclei containing NEB events in cells in each stress condition compared to control cultures grown under normal conditions. Young cells were also examined after separation from old cells and show an increase in NEB compared to the control, but to a lesser extent than old cells. Cells exposed to H_2_O_2_ were grown in different media than the other cultures, and therefore were compared to a different control culture. n is equal to the number of examined sections. (**e**) Thin section containing a NEB event in cells treated with AZC, a chemical which causes proteotoxic stress. (**f**) Percentage of nuclei containing NEB events before as well as 30 and 90 minutes after treatment with AZC. Cells exhibited a higher percentage of NEB (22%) after 90 minutes of exposure to AZC than in any other condition tested. All scale bars 200 nm. Abbreviations: NEB, nuclear envelope budding; N, nucleus; black arrows indicate NEB events. \**P*<.05, \*\**P*<.01, \*\*\**P*<.001 vs. control; ns no significant differences between groups.

Cells of *S. cerevisiae* were also subjected to 90 minutes 0.6 mM H_2_O_2_ (hydrogen peroxide, oxidative stress) and 60 minutes 0.5 mM NaAsO_2_ (sodium arsenite, heavy metal stress). Treated cells, as well as a control cultures for each condition, were cryo-immobilized and prepared for analysis by electron microscopy. Cells that were treated with sodium arsenite had a NEB frequency of 6% (n=249 sections) and cells treated with hydrogen peroxide had a frequency of 5.9% (n=204 sections) (Figure 2**b**-**d**). Both control samples had lower levels of NEB events, as found in undisturbed *S. cerevisiae* (2.4% for the sodium arsenite control for n= 337 and 1.5% for the H_2_O_2_ control for n= 197, Figure 2d). From this set of observations, we conclude that NEB is increased in hydrogen peroxide and sodium arsenite stress as well..

### NEB frequency is highest under induction of protein misfolding

All of the stressors examined above have previously been shown to increase rates of aggregation of misfolded proteins in the cytoplasm and within the nucleus (35, 57–59). To investigate whether or not the observed increase in NEB events is indeed correlated with proteotoxic stress, we examined *S. cerevisiae* cells after treatment with azetidine-2-carboxylic acid (AZC). AZC competes with the amino acid proline during translation and may be mistakenly incorporated into proteins. Since AZC has one fewer carbon atoms in its ring (4 carbon atoms instead of 5 as in proline), its incorporation into a polypeptide causes a different conformation of the amino acid backbone and this results in aggregation of proteins with non-native structures (60, 61). Cells were grown at normal temperature (30°C) and were treated with AZC, cryo-immobilized after 30 and 90 minutes and prepared for analysis by electron microscopy. Cellular viability upon treatment with all of the stressors mentioned above was not compromised, as validated with propidium iodide (PI) staining, indicative of plasma membrane rupture and thus cell death (Figure S3).

The NEB frequency was similar in untreated cells and cells after 30 min of AZC exposure (2.4% n= 337 sections and 2.6% n= 114 sections respectively, Figure 2**f**). However, after 90 minutes of exposure to AZC, the number of NEB events dramatically increased in frequency, reaching 22% (Figure 2**e**-**f**; n= 100). This time-dependent increase in NEB after AZC exposure demonstrates a direct link between the NEB pathway and how the cell accommodates proteotoxic stress.

### NEB events are ubiquitinated

Most proteins that are destined for degradation through the 26S proteasome system must first bind to a poly-ubiquitin chain (62). Moreover, ubiquitin signaling is known to target cytosolic proteins and organelles for degradation by autophagy (63, 64). To probe if the NEB cargo is targeted for protein degradation via ubiquitin signaling, we performed immuno-EM analysis on *S. cerevisiae* cells with a polyclonal antibody that has a stronger affinity to poly-ubiquitin chains than to monomeric ubiquitin (Abcam ab19247). In order to stimulate an increase of the NEB events, cells were subjected to 30 minutes of heat shock. Images of 60 randomly chosen cell sections were recorded and the number of gold particles per area of various cell structures were quantified (Figure 3**a**). Lipid droplets were used as a negative control (4 gold particles/μm^2^) (Figure 3**a**-**b**) and autophagosomes were chosen as a positive control (67 gold particles/μm^2^). The nucleoplasm (excluding areas containing EDC) did not differ from the negative control (4 gold/μm^2^) whereas EDC (protein aggregates) were labeled fivefold more frequently (21 gold/μm^2^) (Figure 3**a**-**b**). Similarly, NEB events (17 gold/μm^2^) were labeled fourfold more frequently than lipid droplets (4 gold/μm^2^). In order to achieve a higher n number for the NEB events, cells were actively traced and selected based on the presence of NEB events. Most of the gold particles identified as associated with NEB events had a preference of localizing in close proximity to the membrane of the bud, but in other examples it would localize more centrally. (Figure 3**b**, **S3**). These results confirm the presence of ubiquitin in both EDC and the NEB events recorded during heat shock in *S. cerevisiae*.

**Figure 3.**
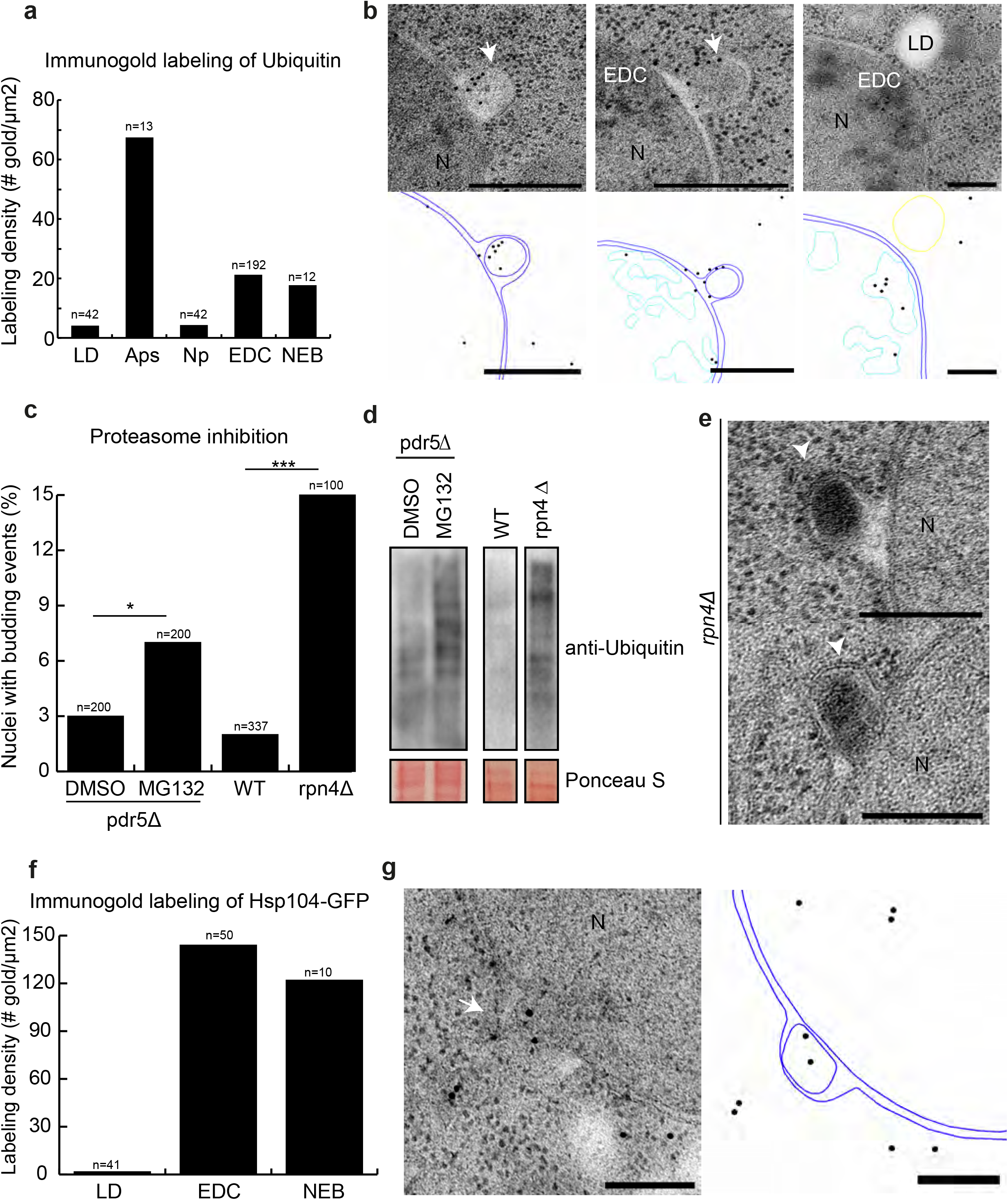
Association of NEB pathway and protein quality control system.. (**a**) Immuno-gold labeling with an anti-ubiquitin antibody (ab19247) was quantified as the amount of gold particles per square micrometer for five different cellular structures. Lipid droplets (n=42) served as a negative and autophagosomes (n=13) as a positive control. EDC (n=192) and NEB events (n=12) exhibited an increase in labeling compared to lipid droplets and the nucleoplasm (n=42) which was defined as area within the nucleus not containing EDC. (**b**) Thin sections labeled with the anti-ubiquitin antibody ab19247. To the bottom of each micrograph a model is drawn to help distinguish gold particles. Two labeled NEB events are shown in the first two panels and an EDC containing gold particles is shown in the third panel. (**c-e**) Proteasomal inhibition triggered a significant increase in NEB events with 73% of them having a distinct electron dense appearance. (**f-g**) Immuno-gold labeling of an anti-GFP antibody of an Hsp104-GFP strain was quantified as the amount of gold particles per square micrometer. Lipid droplets (negative control; n=41) and EDC (positive control; n=50). NEB events (n=10) exhibited an increase labeling density in comparison to the negative control. All scale bars 200 nm. Abbreviations: NEB, nuclear envelope budding; N, nucleoplasm; EDC; electron dense content; LD, lipid droplet; APS, autophagosomes; white arrow indicates NEB.

### Proteasome inhibition increases NEB event frequency

If our hypothesis were correct that NEB could function to remove misfolded proteins from the nucleus that cannot be accommodated by the proteasome, then the inhibition of the proteasome should alter the NEB event frequency. To investigate this, the proteasome was inhibited using two different methods. First, we treated *pdr5Δ* cells (65), deficient in a drug efflux pump with the MG132 drug which is known to partially inhibit proteasomal activity (66). This treatment led to an observed NEB frequency of 7% (n=200), a significant increase compared to the control group (2.5%, n=200) (Figure 3**c**). In order to achieve a stronger proteasomal inhibition, the Rpn4 transcriptional activator of genes encoding proteasomal subunits (43, 67) was deleted and the derived strain was prepared for analysis. Inhibition of the proteasome was confirmed by performing western blots against ubiquitin, since decreased proteasome activity will result in an increase of ubiquitinated proteins (Figure 3**d**). Most of the observed NEB events (11 out of 15) in the *rpn4Δ* cells had a distinct electron dense appearance and no obvious membrane surrounded this electron dense material (Figure 3**e**). In comparison to wild type cells which displayed a NEB frequency of 2.4% (n=337), *rpn4Δ* cells had NEB events in 15% of the sections (n=100) (Figure 3**c**). These results suggest that NEB pathway is an alternative to the ubiquitin proteasome system.

### NEB structures contain Hsp104-GFP

To probe for the presence of aggregated proteins within NEB, we performed immuno-EM in heat shocked cells (38°C for 30 minutes) with an anti-GFP antibody in cells expressing Hsp104-GFP. Even though Hsp104 is not an aggregate per se, it is a well-characterized chaperone which localizes to aggregated proteins in the nucleus and the cytoplasm, where it acts as a disaggregase, (68–70) and supports the degradation of ubiquitinated membrane proteins in the ER-associated degradation pathway(71). In comparison to the negative control (lipid droplets, 1.7 gold/ μm^2^), NEB events were highly labeled by gold particles (122 gold/μm^2^) comparable to the positive control (EDC, 144 gold/μm^2^) (Figure 3**f-g**). This finding is consistent with either a function of NEB in transporting misfolded nuclear proteins to the cytoplasm, or INM proteins to the ER for degradation.

### NEB structures are not misassembled NPCs

It has been suggested the NEB events are misassembled NPCs (54, 72). When certain key genes required for NPC assembly are deleted, herniations of the nuclear envelope similar to NEB events are observed (73, 74). These types of herniations have a distinct morphology with a ‘neck resembling’ formation between the vesicle and the INM which was revealed to be a defective NPC by subtomogram averaging (75). These NPC-exposing herniae are then degraded by autophagy (75, 76). In addition, NPCs assemble via an inside-out mechanism during which the INM evaginates slightly, although functional NPC assembly does not resemble NEB (77).

To clarify if our observed NEB could also be misassembled NPCs, we performed immuno-EM using an antibody that recognizes four NPC components (Nup62, Nup153, Nup214 and Nup358) on both undisturbed *S. cerevisiae* cultures as well as the aged cells. In the undisturbed culture, the majority (67%) of NPCs observed were labeled by gold particles (positive control; n = 95) (Figure 4**a**, 4**e**) and only 16.7% of lipid droplets (negative control; n=82). Only in one case was a NEB event labeled (16.6%, n=6) (Figure 4**c**) whereas all other NEB events were unlabeled (Figure 4**b**-**c**, 4**e**). In the old yeast cells, 81% of the observed NPCs were labeled by gold particles (positive control; n=100) (Figure 4**d**-**e**) whereas only 8% of lipid droplets were labeled (negative control; n=100) (Figure 4**e**). In contrast, only one atypical NEB event was labelled by a gold particle (10%; n= 10) (Figure 4**e**).

**Figure 4.**
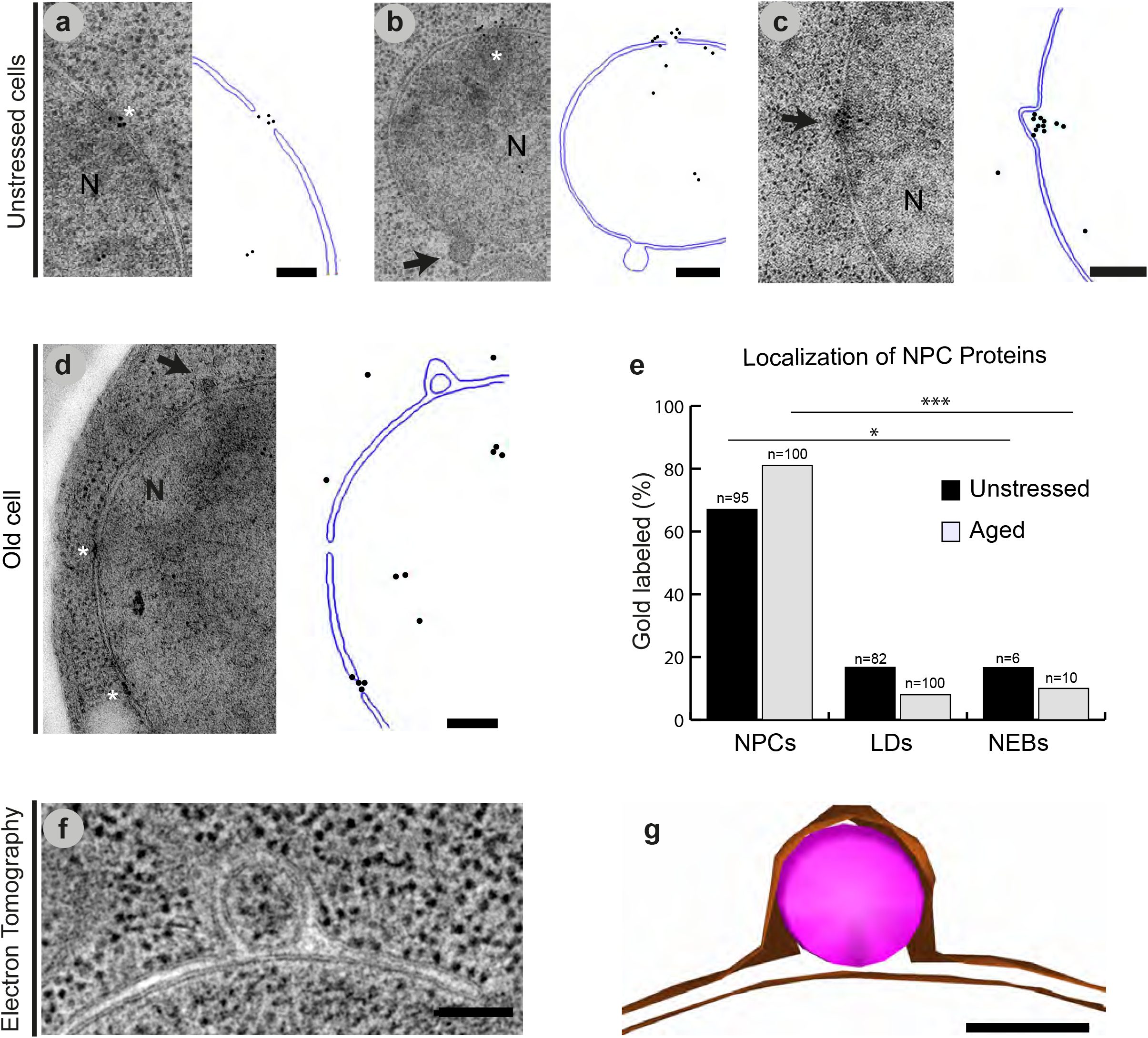
NEB structures are not misassembled NPCs. Immuno-gold labeling, with the Mab414 antibody which recognizes four different NPC proteins, was performed on thin sections of cells grown under normal conditions (**a**-**c**) and old cells (**d**). To the right of each micrograph, a model is drawn to help distinguish the locations of gold particles. (**e**) Percentage of labeled NPCs (positive control), lipid droplets (negative control) and NEB events are plotted. Data for unstressed cells is shown in black, and for old cells in grey. Lipid droplets and NEB events were rarely labeled in contrast to the highly labeled NPCs. (**f**) 25 nm thick slice of NEB in a tomographic reconstruction of *S. cerevisiae*. Both nuclear membranes are clearly visible surrounding a vesicle within the perinuclear space. In all slices of the tomogram, the membrane of this vesicle is complete and not continuous with either nuclear membrane. (**g**) 3D model of the NEB event contained in **f**. Nuclear membranes are presented in orange color and the transporting material in pink. n is equal to the number of examined structures. Scale bars: 200 nm (**a**-**d**), 100 nm (**f**-**g**). Abbreviations: NEB, nuclear envelope budding; N, nucleus; LD, lipid droplet; APS, autophagosomes; NPC, nuclear pore complex; black arrows indicate NEB events, asterisks indicate NPCs. \**P*<.05, \*\*\**P*<.001 vs. NPCs (positive control).

To further support this distinction between NEB and herniation, a comparison of the detailed morphology of NEB events with the previously reported herniations was accomplished through dual-axis electron tomography of the nuclear envelope in *S. cerevisiae*.This analysis showed that the selected NEB event was completely enveloped in a lipid bilayer and wedged in between the two nuclear membranes (Figure 4**f**-**g**). Moreover, the 3D reconstruction revealed that the vesicle contained between the lipid membranes was complete and was not attached to either nuclear membrane by a ‘neck’ with only the outer nuclear membrane extending towards the cytoplasm. As such this morphology was strikingly different to that previously identified as associated with the misassembly of NPCs (77). It is also noteworthy that the 3D reconstruction did not show an obvious connection of the NEB events with the ER, separating these two compartments. A different morphology of a nuclear envelope-ER connection in comparison with the NEB events can also be seen in the acquired thin sections of nuclei (Figure S5). As such, both the immuno-EM results and the 3D morphology of NEB events suggest that the partial or incomplete assembly of NPC and NEB are distinct structures. However, the possibility that a small fraction of the protrusions quantified as NEB events in this study may be better assigned NPC related structures remains.

### The NEB pathway is part of normal cellular function from humans to protists

To gain insights into the evolutionary conservation of NEB we imaged and quantified NEB events in thin sections of nuclei in human mast cells (HMC-1), *C. elegans* nematodes, two yeast species *S. cerevisiae* and *S. pombe*, as well as the parasitic protozoan *T. brucei* using electron microscopy (Figure 5**a**). Since the HMC-1 nucleus has a diameter of ~8 μm (Figure 5**b**) we chose to examine sections corresponding to a little more than one nuclear volume (8 μm/70 nm thick sections ≈ 114 serial sections/nucleus). By the same logic, the number of thin sections examined corresponded to approximately four total nuclear volumes of *T. brucei* and *S. pombe* and five nuclear volumes of *C. elegans* and *S. cerevisiae* cells. For *C. elegans*, images of nuclei were acquired from intestinal cells and oocytes.

**Figure 5.**
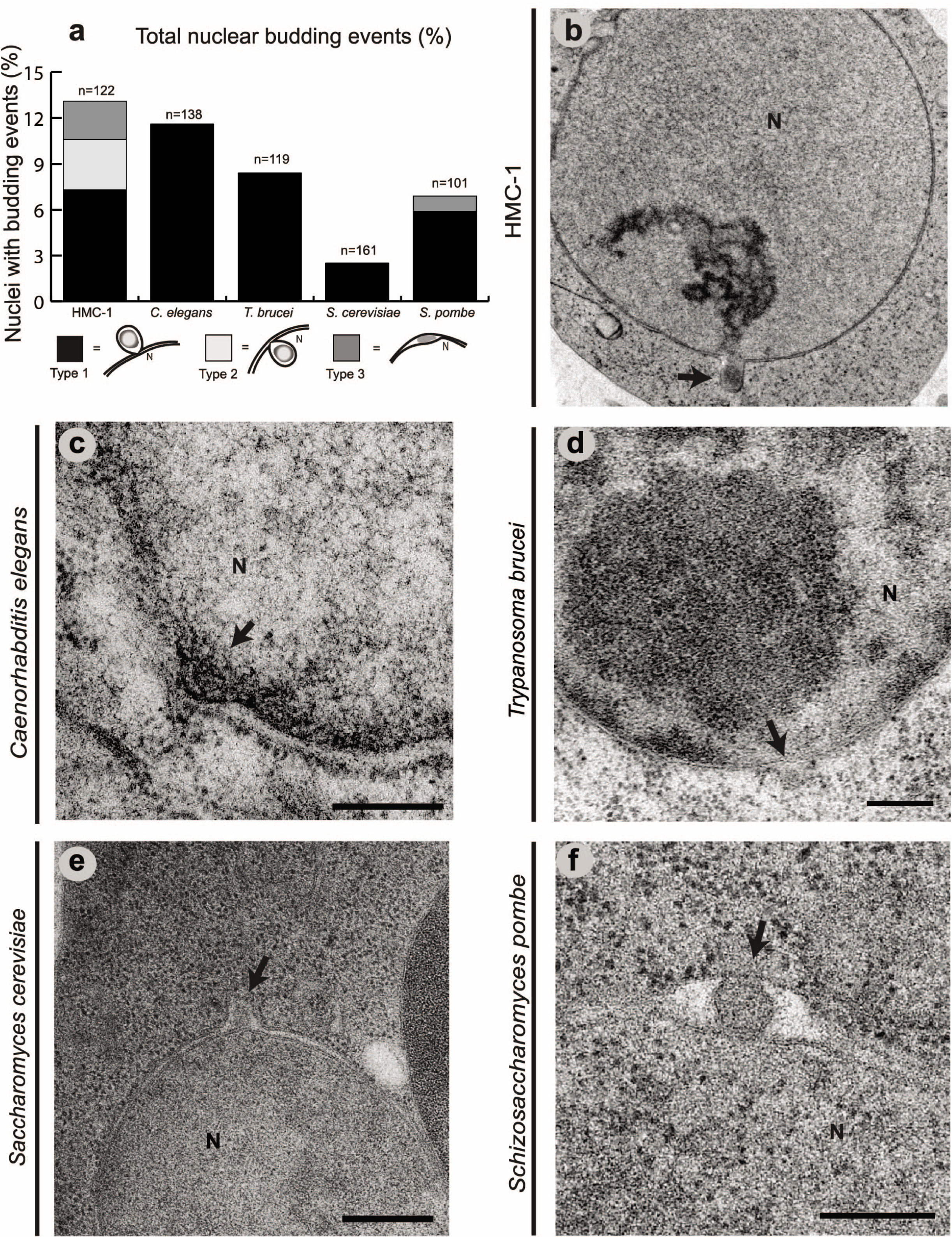
The NEB pathway is part of normal cellular function from humans to protists. Thin sections containing nuclei were examined for the presence of NEB events in five different organisms in wild type cells grown under normal conditions. (**a**) Percentage of nuclei containing NEB events. NEB events were either outwards protruding (type 1), inwards protruding (type 2), or did not protrude more in either direction but clearly contained a particle in the perinuclear space (type 3). NEB was observed in (**b**) HMC-1 cells (n=122) (**c**) *C. elegans* (n=138) (**d**) *T. brucei* (n=119) (**e**) *S. cerevisiae* (n=161) and (**f**) *S. pombe* (n=101) by electron microscopy. Scale bars: 2 μm (**b**), 200 nm (**c**, **f**), 500 nm (**d**), and 300 nm (**e**). Abbreviations: NEB, nuclear envelope budding; HMC-1, human mast cell line 1; N, nucleus; black arrows indicate NEB events.

In HMC-1 cells, three morphologically different categories of NEBs were observed (Figure 5**a**). The most commonly observed morphology was outwards protruding buds, occurring in 7.3% of nuclear sections (n = 9; Figure 5**a**-**b**), which will hereafter be referred to as type 1 NEB. In a few cases of type 1 NEB events, both the outer and inner membranes expand towards the cytoplasm with electron dense material partially wrapped by the nuclear envelope bud and the protrusion is open to the nucleoplasm (Figure 5**b**). In most of the type 1 NEB events however (n= 6), the electron dense material is clearly located between the two membranes with only the outer membrane expanding towards the cytoplasm. The second type of NEB events observed had the inner nuclear envelope expanding towards the nucleoplasm. These were termed type 2 NEB events, and occurred in 3.3% of sections. Finally, for type 3 NEB events the transported material is situated between the nuclear membranes with no clear protrusion in either direction. This occurred in 2.5% of sections. Some thin sections of nuclear envelopes contained more than one bud, revealing that NEB can occur at multiple sites on the same nucleus (Figure S1).

NEB events are observed to occur in all five organisms examined (Figure 5**a**) but with varying frequency. The lowest frequency of NEB events was in *S. cerevisiae* (2.5%, n=161 sections), followed by *S. pombe* (6.9%, n=101 sections), *T. brucei* (8.4%, n=119 sections), *C. elegans* (11.6%, n=138 sections) and most frequently in HMC-1 cells (13.1%, n=122). Only type 1 NEB events were identified in all organisms other than HMC-1 (Figure 5**b**-**e**), except for a single event in *S. pombe* that we classified as type 3 (Figure 5**f**). The morphology of the NEB events varied between and within species, with some buds being small and having indistinct electron dense material whereas other events were large and had a membrane surrounding the material. The average size of the NEB events did not show a large variation between species except for HMC-1 cells, which yielded distinctly larger buds (Figure S6).

The morphology of NEB was compared in more detail through electron tomography reconstructions of NEB events from both *S. pombe* and *T. brucei*. In *S. pombe*, one event of type 1 NEB was visualized (Figure 6**a**-**c**) whereas two type 1 NEB events were reconstructed from *T. brucei* (Figure 6**d**-**h**). The NEB event that appeared to have progressed the furthest had distinct regions of higher electron density surrounding the base of the bud (Figure 6**g**, black arrow). For both *S. pombe and T. brucei* the cargoes of the NEB events were found between, and fully separated from, the nuclear membranes, and were similar to what was observed in *S. cerevisiae* (Figure 4**g**). Moreover, the NEB event observed for *S. pombe* clearly displayed the presence of a lipid bilayer that completely enveloped the cargo (Figure 6**b**). However, a clear bilayer surrounding the transported material could not be ascertained in the *T. brucei* tomographic reconstruction. Since in many cases the transported material appeared to have a vesicle-like structure, we tested for an involvement of the ESCRT pathway known to facilitate the formation of cellular vesicles, and already implicated in the formation of morphologically similar protrusions of the nuclear envelope (78, 79). However, ESCRT components did not appear to contribute to the formation of NEB events in our experiments (S7).

**Figure 6.**
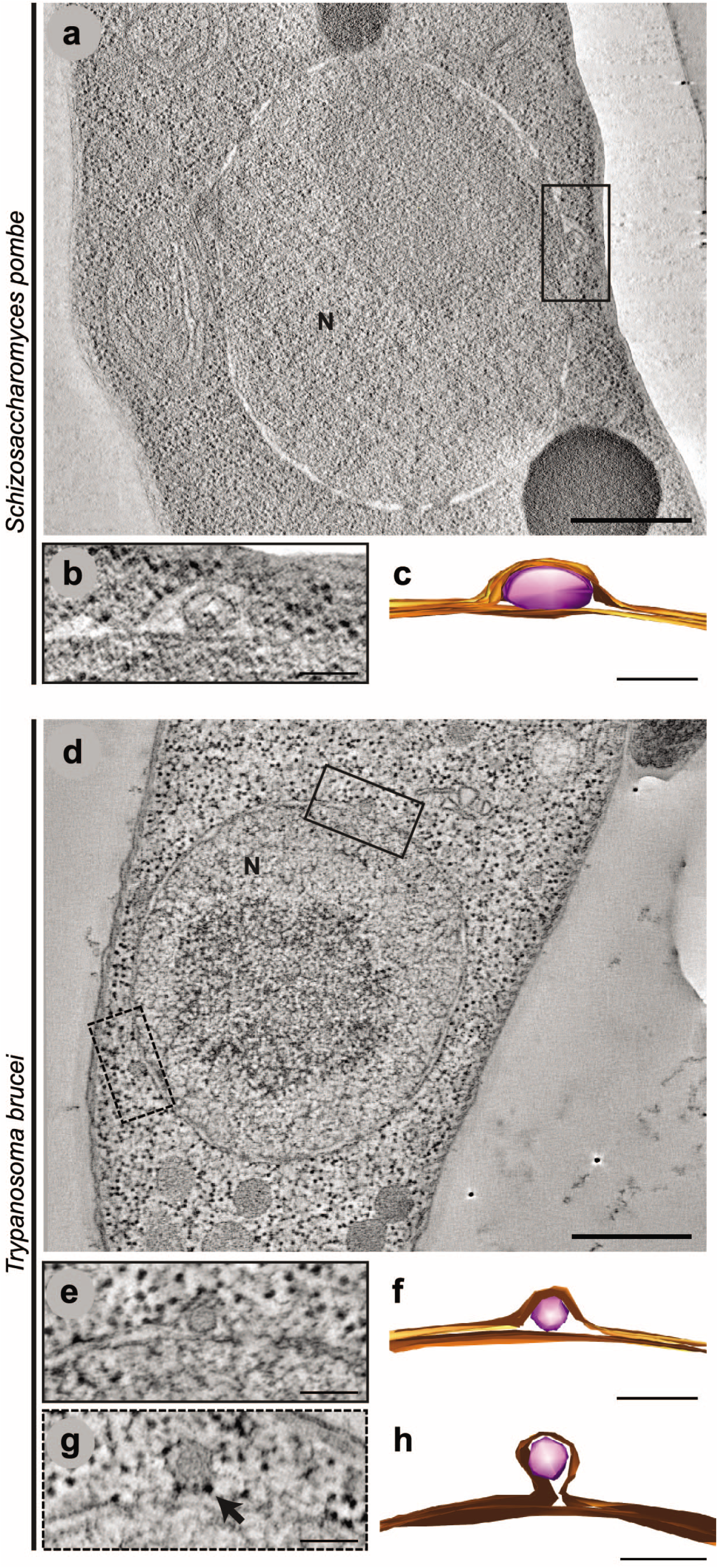
3D architecture of NEB events of *S. pombe* and *T. brucei* visualized with electron tomography. (**a**, **d**) 1.5 nm thick slices of *S. pombe* and *T. brucei* tomographical reconstructions showing NEB events (black boxes). A second NEB event in *T. brucei*, not clearly visible in the presented tomographic tilt, is located on the left lower part of the nucleus (dashed black box). (**b**, **e**) 15 nm thick zoomed-in views of the NEB events in *S. pombe* and *T. brucei*. In *S. pombe* a lipid bilayer is surrounding the transported material whereas in *T. brucei* the density is surrounded by the nuclear membranes only. (**g**) Second NEB event of *T. brucei* (acquired at the tomographic tilt that allowed best visualization of this NEB event). Electron dense particles are observed around the neck of the bud (black arrow). (**c**, **f**, **h**) Reconstructed 3D models of the NEB events. Nuclear membranes are presented in orange color and the transporting material in pink. Scale bars: 500 nm (**a**, **d**) and 100 nm (**b**-**c**, **e**-**h**). Abbreviations: NEB, nuclear envelope budding; N, nucleus;

We conclude that NEB was observed in all species examined in this study. Our findings combined with previous published observations are summarized into a phylogenetic tree (Figure 7), illustrating how the NEB pathway is evolutionarily conserved among eukaryotes and is part of normal cellular function in an evolutionarily diverse sampling of species.

**Figure 7.**
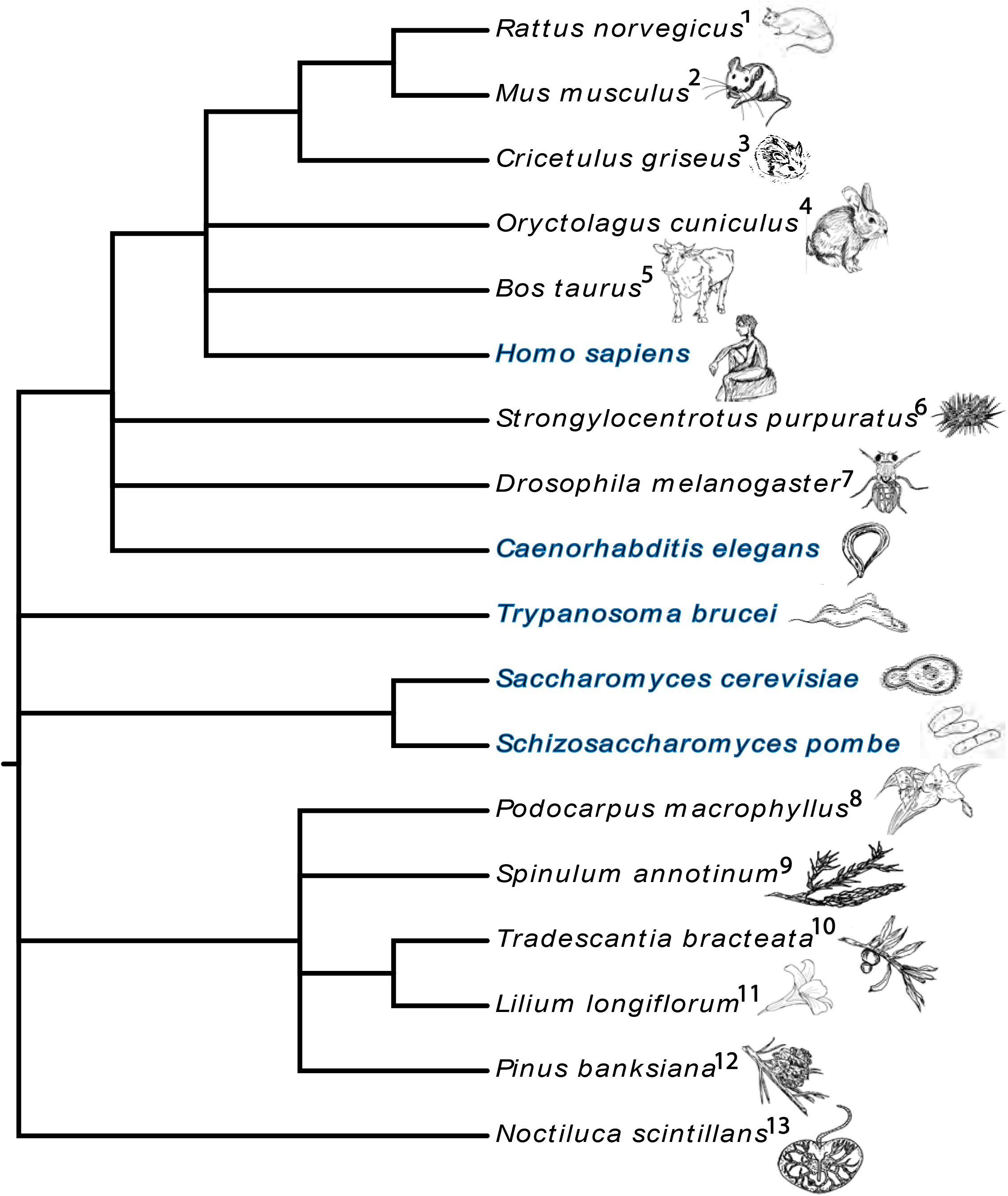
Phylogenetic tree of the species where NEB events have previously been observed. The phylogenetic relationships between the species are presented as a horizontal cladogram with the root to the left. Species indicated with blue color are from this study. References presented from 1 to 13 respectively: (17), (18), (14), (16), (15), (29), (19, 26, 28), (25), (21), (20), (23), (24), (22). All sketches are drawn by the author and are not subject to copyright protection.

## Discussion

Nuclear envelope budding (NEB) has been observed since 1955 as nuclear herniations across a diverse range of organisms and at different developmental stages. This phenomenon is well studied with regards to its role in virus infection and the detailed structure of NEB events have been revealed using cryo-electron tomography of infected cells (11). Yet, NEB remains viewed as an irregularity rather than a normal route for transport over the nuclear envelope in healthy mature cells. We present here three different lines of evidence that NEB is a widely conserved, physiologically normal cellular process that increases in frequency due to cellular stress and is specifically activated by an increase in protein aggregation. Firstly, we observed that five distinct stress conditions: heat shock; hydrogen peroxide; arsenite; proteasome inhibition and AZC treatment, all led to an increase in the frequency of NEB, with the AZC treatment most prominently activating the NEB pathway among all stressors. Secondly, immuno-EM revealed the presence of ubiquitin and Hsp104, a protein disaggregase, in the cargo of the buds, which supports our hypothesis that this pathway is involved in protein degradation. Thirdly, NEB events were detected in every species examined, from T. brucei to human cell line HMC-1. Collectively, these results suggest a role of the NEB pathway in protein quality control and highlight its evolutionary conservation. These observations shed light on the cell’s mechanism to cope with proteotoxic stress and identify a new module of the nuclear quality control system. Moreover, since the NEB pathway is evolutionarily conserved and is also present in human cells, this work provides a new perspective for studying cellular stress response related human diseases (80).

### Nuclear budding and the protein quality control system

Although chaperones and the ubiquitin-proteasome system are known to clear misfolded proteins from the nucleus (81, 82), the nuclear protein quality control system is not as well characterized as the cytosolic and the ER systems (39). Various cellular events and pathways have been described whereby aggregated proteins and chaperones are imported from the cytosol and into the nucleus and these pathways are consistent with observations that 80% of the proteasomes at steady state are located within the nucleus (83, 84). An alternative pathway, however, has also been reported for which ubiquitinated proteins in mammalian cells and in the nematode *C. elegans* are transported from the nucleus to the cytosol via a ubiquitin-associated domain-containing protein (UBIN) (85).

Some proteins that exceed the size limit of the NPCs may nevertheless enter the complex after a reversible conformational change in their tertiary structure (86). Although large aggregates could potentially be transported in a similar manner, it stands to reason that under conditions of cellular stress, the transportation of many aggregates through NEB events would provide a more efficient and therefore advantageous gateway into or out of the nucleus. In support of the above supposition, a process called “piecemeal microautophagy of the nucleus” has been described in *S. cerevisiae* in which nuclear envelope buds are released directly into the vacuole lumen for degradation (49).

Our observations that NEB increases during five different cellular stresses (heat shock, hydrogen peroxide, arsenite, proteasome inhibition and AZC exposure) in *S. cerevisiae* strongly implicate that NEB is related to the protein quality control system. Aging has been previously shown to influence the formation of NEB-like events in the perinuclear space (54), with an apparent increased in replicatively aged cells of about 17% when compared to the remaining young population (stated as mixed population). Though the isolation method we used in this study involved a higher number of isolation steps in comparison with the aforementioned literature, we were unable to reproduce the difference in NEB frequency between the old and young population. There is a significant increase in both isolated cell groups with a stronger effect on the aged cells. Possible reasons for the inability to reproduce what has been shown before could be due to the differences in the isolation process. Conclusions that can be safely extracted are that mechanical stress is also able to increase NEB frequency and aging most likely creates a further stressful environment for the cells.

The presence of ubiquitin and Hsp104 in the NEB events lends further support for this functionality due to the fact that ubiquitin is involved in many protein quality control pathways including the ubiquitin-proteasome system and macroautophagy, and Hsp104 is known to mediate disaggregation of misfolded proteins (87–89). The large increase in NEB events due to the AZC treatment reinforces this putative role of NEB as a cellular stress response. Since AZC acts upon protein conformational stability directly without influencing other parameters such as the external temperature or membrane fluidity, these experiments constitute a more targeted challenge for the protein quality control system. Additionally, in a previous study where aggregated proteins were identified after cellular stress induced by AZC, arsenite and hydrogen peroxide treatment, AZC had the largest number of affected proteins among the stressors (58). A strong effect of AZC on the folding of proteins could explain why NEB is more prominent in the AZC-induced stress response when compared to the cellular response to other stressors. Finally, the connection between proteasomal activity and NEB frequency supports the association with the protein quality control system. Inhibition of proteasomal activity showed a clear increase in NEB events which could be translated as a cellular attempt to cope with a large number of misfolded proteins or proteins which are normally degraded by the proteasome. Most of the NEB events recorded in the *rpn4Δ* strain, had a distinct morphology compared to NEB events triggered by other stressors. This observation may reflect a difference in the nature and composition of the transported material, indicating that NEB may well be involved in more than one cellular activity.

### The distinction between NEB events and defective NPCs

Deleting key genes required for NPC assembly can induce nuclear envelope “herniations” that appear somewhat similar in structure to NEB events (73–76, 90, 91). These herniations are believed to arise from the outer nuclear membrane sealing above the defective NPC, thereby capturing the transported material between the NPC and the nuclear envelope and creating a bulge resembling a budding vesicle. If this phenomenon underpinned our observed NEB events, the intermembrane vesicle should be connected to the INM where the defective NPC is located. A connection between the INM and intermembrane vesicle resembling a “neck” would also be a plausible intermediate state in the formation of a vesicle from the INM that encloses nuclear material. Such neck-like connections between the INM and contents of the bud were previously observed in strains deficient in TorA, an ER protein with ATPase activity, and various Nups that lead to defective NPCs (31, 73, 74). Furthermore, the size of the necks seen in NEB events after hyperactivation of Chm7, a component of the ESCRT-III like complex, and misassembled NPCs were compared previously and it was concluded that while appearing similar, both features were morphologically distinct (79). Neck-like or NPC-like structures connecting the intermembrane vesicles to the INM were not visible in any of our acquired tomograms, suggesting that our tomograms contain buds in which the intermembrane vesicle or material has fully formed and these buds are not the result of NPC malformation. In addition to these morphological differences revealed by 3D tomographic reconstructions, the immuno-EM assay demonstrated that the NEB events were not associated with NPCs since only two of the 16 observed events in this assay were labeled for NPC proteins.

### The evolutionarily conserved nature of NEB

We observed NEB events within the cells of all species we examined including animals, fungi and protozoa. There is a thread stretching back through the scientific literature which shows that NEB events have repeatedly been observed in animals, plants and protists (Table S1). The genetic diversity of these observations is summarized in Figure 7, which presents a phylogenetic tree illustrating that NEB events have been observed for 18 different species within the eukaryotic domain. Therefore, despite NEB not being widely recognized as a means of nuclear transport, these observations combine to create a compelling argument that NEB is an evolutionarily conserved phenomenon of eukaryotic cells.

From our sequence of experiments using the model systems *S. cerevisiae* we concluded that NEB events facilitate the removal of aggregated proteins from the nucleus. This pathway could potentially also support the co-transportation of other cellular components required for protein degradation. Since NEB embodies an alternative route for transport over the nuclear envelope that has been largely overlooked, our observations highlight an opportunity for discovery in eukaryotic cell biology where a broad range of interconnected questions may emerge. It will be necessary to determine if the frequency of NEB is also increased by cellular stressors in human cells, and in particular additional cellular stress associated with aging. One issue is the important role of protein aggregation within a variety of neurodegenerative diseases such as Alzheimer’s and ALS (80, 92). Should NEB also be an underappreciated mechanism for the clearance of protein aggregates in human neurons then this potentially has far-reaching neurological implications.

## Materials and Methods

### Antibodies and reagents

Please refer to Supplementary tables 3 and 4 for a detailed information on antibodies, strains, primers and reagents.

### Experimental models

#### HMC-1 cells

Human mast cell line 1 (HMC-1; (93)) cells were cultured in Iscove’s modified Dulbecco’s medium (IMDM). The cells were then treated with 10% exosome-depleted fetal bovine serum (FBS), 100 units/ml streptomycin, 100 units/ml penicillin, 2 mM L-glutamine, and 1.2 U/ml alpha-thioglycerol in incubators kept at 37°C and 5% carbon dioxide (94, 95).

#### Trypanosoma brucei

Procyclic *T. brucei* strain 427 was cultured in SDM-79 media with 20% fetal bovine serum. Cultures were prepared and maintained at a concentration of 5 × 10^5^ and 1 × 10^7^ cells per mL (96–98).

#### Schizosaccharomyces pombe

Logarithmically growing wild type fission yeast *S. pombe* were grown at 30°C in YE5S medium (94, 99, 100).

#### Caenorhabditis elegans

The *C. elegans* wild type reference strain was the Bristol N2 variety. The worms were cultured on normal growing media plates (NGM plates) and the *E. coli* strain OP50 was used as a food source. The worms were maintained at the optimal temperature of 25°C (101, 102). Adult worms were used for electron microscopy.

#### Saccharomyces cerevisiae

Wild type cells of *S. cerevisiae* (BY4741) were cultured in YPD media at 30°C (103). A strain with endogenous *HSP104* C-terminally tagged with GFP (104) was used for old cell isolation, heat shock and sodium arsenite experiments. The deletion mutants *hsp104Δ* and *rpn4Δ* are from the YKO collection (EUROSCARF, Frankfurt, Germany). Strains used for investigating involvement of the ESCRT-pathway in NEB and the *pdr5Δ* strain are from this study (see yeast strains list). Transformations were performed following standard procedures (105), and gene deletions and endogenous tags were integrated via homologous recombination (106). For the hydrogen peroxide experiment, cells were grown in synthetic complete media (with yeast nitrogen base, without amino acids, pH 5.5, complete supplement mixture of amino acids and 2% glucose).

### High-pressure freezing for electron microscopy and tomography

All samples used in this study has been prepared using high-pressure freezing followed by freeze substitution. HMC-1 cells were loaded into membrane carriers and were high-pressure frozen in a Leica EM PACT1 (Leica microsystems, Wetzlar, Germany). *T. brucei* and *S. pombe* were loaded into carriers and were high-pressure frozen in a Leica EM PACT2. *C. elegans* and *S. cerevisiae* samples were loaded into aluminum specimen carriers and were high-pressure frozen in a Wohlwend Compact 3 (M. Wohlwend GmbH, Sennwald, Switzerland). Yeast paste from wild type *S. cerevisiae* cells was used as a cryoprotectant filling the space surrounding the worms. For a summary of all high-pressure freezing and freeze substitution experiments see table 1. For all samples (except *S. pombe*) a short freeze substitution protocol was applied, using 2% uranyl acetate dissolved into acetone (UA; from 20% UA stock in methanol) for one hour (94, 97, 107). To increase the penetration into intact worms, the UA solution was left on the samples for 14h (as the temperature was increased to −50°C). The UA incubation was followed by two washes in 100% acetone for one hour each. Before embedding, the temperature was raised from −90°C to −50°C overnight with a rate of 3°C/h. Samples were then embedded in K4M or HM20 resin in increasing concentrations of 20%, 40%, 50%, 80% and finally three times in 100% plastic (2 hours per solution). Polymerization of the plastic occurred over 48h using UV light at −50°C followed by 48h in room temperature. *S. pombe* cells were high-pressure frozen and fixed through freeze substitution (long protocol) with anhydrous acetone containing 0.25% UA, 0.1% dehydrated glutaraldehyde and 0.01% osmium tetroxide (OsO_4_) (99, 100, 108, 109). For tomography, serial semi thick sections of about 210-250 nm (*S. pombe*) and 350 nm (*T. brucei* and *S. cerevisiae*) were cut and samples were poststained with 2% UA in dH_2_O followed by Reyonold’s lead citrate. Gold particles (15 nm) from British Bio Cell were applied to both sides of the grid to be used as fiducial markers. (96, 97, 99).

All other samples were sectioned in 70 nm thin sections and placed on either copper slot or mesh grids. Sections were stained with 2% UA for 5 minutes and Reynold’s lead citrate for 1 minute (110). Washing steps after each staining were performed in dH_2_O.

### Immuno-electron microscopy

For the immuno-labeling experiments, the same high-pressure frozen samples embedded in HM20 resin were used, a benefit of that short FS protocol (107). For a summary of these experiments, see table 2. Grids with 70 nm thick sections were fixed in 1% paraformaldehyde (PFA) in PBS for 10 minutes. After three PBS washes of 1 minute each, samples were blocked with 0.1% fish skin gelatin and 0.8% BSA in PBS for 1 hour. For detection of NPC proteins, grids were then incubated in a 1:50 dilution of mAb414 (BioLegend, San Diego, USA) for two hours, followed by a 1:150 dilution of rabbit anti-mouse immunoglobulins (Agilent/Dako, Glostrup, Denmark) for an hour, and then a 1:70 dilution of 10 nm gold-conjugated protein A (CMC UMC Utrecht, The Netherlands) for 30 minutes. For labeling of ubiquitin, grids were incubated in a 1:20 dilution of antibody ab19247 (Abcam, Cambridge, UK) for 2 hours. Detection of GFP was performed by using a 1:5, 1:10, or 1:30 dilution of ab6556 (abcam, Camebridge, UK) and detection of Hsp104 by using a 1:100 dilution of ab69549 (abcam, Camebridge, UK) incubated overnight. Goat-anti-Rabbit IgG 10 nm gold (Electron Microscopy Sciences, Hatfield PA, USA) was then used at a 1:20 dilution for an hour. All incubations were performed at room temperature, except for the primary antibody which was kept at 4°C. Three washing steps (20 min in PBS) were carried out after incubations with each antibody. 2.5% glutaraldehyde was applied to sections for 1 hour followed by three washes (1 min in dH_2_O). Sections were then contrast stained in 2% UA for 5 minutes (wash 3x 2 min in dH_2_O) and 1 minute in Reynold’s lead citrate (washed 5x 1 min in dH_2_O).

### Image acquisition and electron tomography

All thin sections were imaged at 120 kV either on a LEO 912 OMEGA (Zeiss, Krefeldt, Germany) equipped with a 2k x 2k VELETA Olympus CCD camera or on a Tecnai T12 transmission electron microscope equipped with a Ceta CMOS 16M camera (FEI Co., Eindhoven, The Netherlands). Double axis tilt series of serial sections of *T. brucei* samples were acquired every degree using the serialEM software (111) on a Tecnai TF30 300 kV IVEM microscope (FEI Co., The Netherlands) equipped with a Ultrascan 785 4k x 4k camera binned to 2k x 2k (pixel size 1 nm). For *S. pombe*, digital images (Gatan Ultrascan 890 or 895 camera, pixel size 1.5 nm) single axis tilt series were taken every 1.5° over a ±60°−65° range operating a Tecnai TF20 electron microscope (FEI Co., Eindhoven, The Netherlands) (99). For *S. cerevisiae*, double axis tilt series of serial sections were acquired on a Tecnai TF30 300 kV microscope (FEI Co., The Netherlands) equipped with a Gatan One View camera (pixel size 1.6 nm, increment 1.5° over a ±60° range). Tomograms were acquired with the use of serialEM (111). The tomographic reconstruction was performed using of the IMOD software package (112).

### Isolation of old yeast cells

Isolation of old yeast cells was performed according to Smeal et al. with some modifications in order to reduce mechanical stress and improve the cell morphology for electron microscopy (56). In brief, the cell surface of exponentially growing cells was labeled with biotin by incubating the cells with 0.5 mg/ml Sulfo-NHS-LC biotin (#21335, Thermo Fisher scientific) for 20 min at room temperature. Cells were grown in YPD, harvested prior saturation of culture, and washed in PBS + 0.5% glucose. Biotinylated cells were labeled with 17.5 ug/ml Streptavidin magnetic beads (#21344, Thermo Fisher scientific) for 1.5 hours followed by 3 x 15 min magnetic sorting with PBS + 0.5% glucose washes in between. Another two rounds of growth, streptavidin-labeling, and sorting was performed. The old cells and their unbound daughters were recovered in YPD for 4 hours before high-pressure freezing.

### Stress treatments of *S. cerevisiae*

Cells were grown at 30°C to mid exponential phase. For mild heat shock, the culture was shifted to 38°C and samples were collected after 0, 5, 15, 30, 45, and 90 minutes (113). The more severe heat shock was performed at 42°C for 30 min. For hydrogen peroxide, sodium arsenite, and AZC treatments the culture was split into two, one kept as unstressed control and one treated with the stressor. Cells were exposed to 0.6 mM hydrogen peroxide for 90 minutes, 0.5 mM sodium arsenite for 60 minutes, or 1 mg/mL AZC for 30 and 90 minutes (corresponding control culture was grown for 60 minutes after mid exponential phase). Cells were harvested by filtration followed by high-pressure freezing.

### Proteasome inhibition

The *rpn4Δ* strain and a wild type control was grown to mid exponential phase and prepared for high-pressure freezing. The *pdr5Δ* strain was grown to mid exponential phase prior addition of either 50 μM MG132 (#C2211, Sigma-Aldrich) dissolved in DMSO or an equivalent volume of DMSO. A 60 min incubation at 30°C followed before cells were prepared for high-pressure freezing.

### Analysis of polyubiquitination

Cells were cultured and treated as described above. Mid-exponentially grown yeast corresponding to OD_600_ = 20 was harvested, washed once in distilled water and resuspended in 1 ml lysis buffer (100 mM Tris pH 7.5, 100 mM NaCl, 5 mM ethylenediaminetetraacetic acid, 1 mM dithiothreitol, 1 mM phenylmethylsulfonyl fluoride, 20 mM N-ethylmaleimide). Homogenisation was conducted via an Avestin Emulsiflex C-15, applying a homogenization pressure of 18 000 psi. Samples were cleared from cell debris and unlysed cells by centrifugation for 5 min at 3500 rcf and 4°C. 100 μl of cleared lysate was mixed with the same volume of 2x Laemmli buffer (100 mM Tris pH 6.8, 4% SDS, 20% glycerol, 0.2% bromophenol blue, 200 mM 2-mercaptoethanol) and incubated for 15 min at 95°C. The samples were then applied for SDS-PAGE and immunoblotting following standard protocols. Equal loading was controlled via Ponceau S staining directly after wet electrotransfer on PVDF membranes and blots were decorated with an anti-ubiquitin antibody (1:1000, HRP-conjugated, P4D1, sc8017, Santa Cruz). Clarity Western ECL Substrate (BIO-RAD, 1705060) and a ChemiDoc XRS+ Imaging System (BIO-RAD, 1708265) were used for detection

### Analysis of cell death

Loss of membrane integrity was assessed with propidium iodide (PI) staining as previously described (114). Briefly, cells were harvested in 96-well plates after the respective stressor/mock treatment and resuspended in 250 μl of PI solution (100 μg/l PI in phosphate buffered saline PBS; 25 mM potassium phosphate, 0.9% NaCl; adjusted to pH 7.2). After 10 min incubation in the dark, cells were washed with 250 μl of PBS and analysed via flow cytometry (Guava easyCyte 5HT; Merck group). 5000 events were recorded per strain and condition using InCyte software (3.1).

### Confocal microscopy

For visualization of nuclei, yeast cells were harvested and resuspended in DRAQ5 staining solution (5 μM DRAQ5 in PBS). After 10 min incubation in the dark, cells were washed once with PBS and immobilized on agar slides. Specimen were analyzed with a ZEISS LSM700 microscope using ZEISS ZEN blue software control. Plan-Apochromat 63x/1.40 Oil M27 objective was employed. Appropriate filter settings were used to visualize GFP and DRAQ5. Micrographs were analyzed and processed with the open-source software Fiji (115). To reduce image noise, Gaussian filtering (σ = 1) was applied, followed by background subtraction (rolling ball radius = 100 pixels). Pictures within an experiment were captured and processed in the same way.

### Phylogenetic tree

A phylogenetic tree was constructed showing all the organisms where NEB events have been observed. The tree was generated based on the NCBI taxonomy browser for scientific names and visualized by the EvolView online software (http://www.evolgenius.info/evolview/).

### Statistics and reproducibility

The frequency of NEB events was achieved by counting the number of events present in approximately 100-200 thin sections of nuclei. The percentage of sections containing events in each organism was presented as a bar graph. For the NPC immuno-EM assay, different cell compartments (NPCs, NEBs and lipid droplets) were categorized as labeled or not labeled based on the presence or absence of gold particles. Similarly, the percentage of labeled compartments was presented as a bar graph. For the statistical analysis, different statistical tests were examined but a non-parametric Wilcoxon test was performed as our data represent frequencies and the numerical values are not following a normal distribution nor are continuous variants (values were considered as 0 for the absence of events and 1 for the presence of events).. The test was performed using the MATLAB multi-paradigm programming language. For further details on the examined statistic tests see Supplementary table 2.. For the ubiquitin and Hsp104-GFP immuno-EM, the area and number of gold particles of different cell compartments were measured using IMOD (https://bio3d.colorado.edu/imod/). The amount of gold particles per area was presented as bar graph. For cell death analysis, results are displayed either as dot plots (if n ≤ 5) where mean (square), median (centre line) and s.e.m. are depicted, or as box plots (if n > 5) with mean (square) and median (centre line) as well as whiskers presenting minima and maxima within 2.2 interquartile range (IQR). Outliers were defined by the 2.2-fold IQR labelling rule. Normal distribution of data was examined with a Shapiro-Wilk’s test and homogeneity of variances was evaluated with a Levene’s test. A student’s t-test was used to compare between two groups and displayed significances are two-sided, and a one-way ANOVA with a Bonferroni post hoc test was applied to compare between three and more groups. Significances are indicated with asterisks: \*\*\*\**p*<.0001, \*\*\**p*< .001, \*\**p* < .01, *\*p* < .05, n.s. not significant. Further information on sample size can be found in the respective figure legends. All figures were processed with Origin Pro 2017 or Adobe Illustrator CS6 (Adobe).

## Data availability

Links to the online library for the 2D electron microscopy pictures acquired during this study:

http://cellimagelibrary.org/groups/50813

http://cellimagelibrary.org/images/50818

http://cellimagelibrary.org/groups/50832

http://cellimagelibrary.org/groups/50841

http://cellimagelibrary.org/groups/50849

http://cellimagelibrary.org/groups/50873

http://cellimagelibrary.org/images/50882

http://cellimagelibrary.org/images/50888

http://cellimagelibrary.org/images/50900

http://cellimagelibrary.org/groups/50906

http://cellimagelibrary.org/images/50920

http://cellimagelibrary.org/groups/50935

http://cellimagelibrary.org/images/50945

## Summarized protocols

Please refer to supplementary table 5 for a summarized information on thin sections, tomograohy samples and immuno-EM samples preparation.

## Acknowledgments

This project was funded by the Swedish Research Council Grant (2015-00560) and the Knut and Alice Wallenberg Foundation (grant KAW 2012.0284, KAW 2014.0275) to RN, the Swedish Research Council Grants (621-2014-4597, 2018-03577) to MJT, the Swedish Research Council Young Investigator Grant (2015-05427), the Austrian Science Fund (J4342-B21) to VK and Swedish Research Council grant (2019-04004) to JLH and Knut and Alice Wallenberg grant (KAW 2017.0091) to TN and JLH.

All authors would like to thank Marc Pilon for providing the *C. elegans* worms and for contributing intellectually to the manuscript writing. Further on, we would like to thank Rebecca Andersson for her help in conducting the hydrogen peroxide experiment and Per Widlund for reagents. We would also like to thank Claes Andreasson, Per Widlund and Martin Ott for helpful discussions.

## Author Contributions

Conceptualization, D.P., J.T.C., S.B. and J.L.H.; Methodology, D.P., J.T.C., L.L.B., K.K. and J.L.H.; Validation, D.P., J.T.C. and K.K.; Formal Analysis, D.P., J.T.C., R.N. and K.K.; Investigation, D.P., J.T.C., K.K., L.L.B., V.K. and S.A.; Resources, S.B., M.J.T., T.N., R.N. and J.L.H.; Writing-Original Draft, D.P., J.T.C. and J.L.H.; Writing-Review & Editing, D.P., J.T.C., K.K., L.L.B., S.A., V.K., S.B., M.J.T., T.N., R.N. and J.L.H.; Visualization, D.P., J.T.C. and J.L.H.; Supervision, L.L.B., R.N., M.J.T., T.N., S.B. and J.L.H.; Funding Acquisition, V.K., S.B., T.N., R.N., M.J.T. and J.L.H.

