## Supplemental info for "Nuclear envelope budding is a response to cellular stress"

#### Supplementary figure 1 (Related to Figure 5)

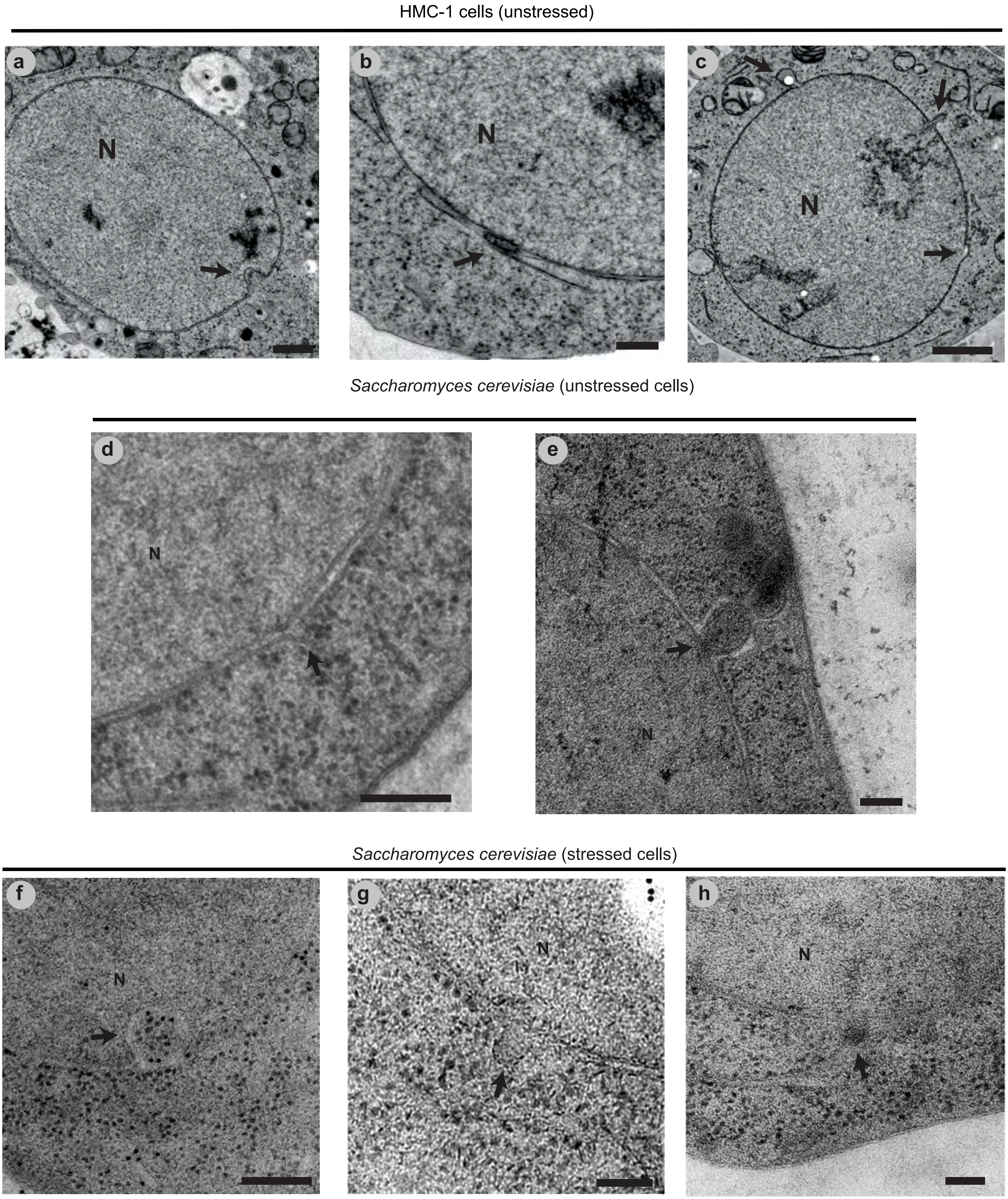

**Supplementary figure 1. Different morphologies of NEB events.** Electron microscopy of thin sections revealed several NEB events in (**a-c**) HMC-1 cells and (**d-h**) *S. cerevisiae* with unique morphologies. These could correspond to different stages in the same process, or a variety of processes that occur by NEB. (**a**) NEB in which the event protruded towards the nucleoplasm (Type 2 NEB). (**b**) NEB in which a vesicle was clearly inside the perinuclear space but with no distinct directionality (Type 3 NEB). (**c**) Two NEB events occurring in the same nuclear section. (**d**) NEB event showing only a small protrusion of the outer nuclear membrane with no material apparent inside. (**e**) Outwards protruding NEB event containing two vesicles instead of one. One vesicle appears complete however the second is still continuous with the inner nuclear membrane. (**f**) Inwards protruding NEB observed in a yeast culture that was subjected to heat shock. The event contains a complete vesicle within, and the contents have similar electron density to cytoplasm with ribosomes clearly visible. (**g**) NEB event showing only a small protrusion of the outer nuclear membrane and no material apparent inside. This event was observed in an aged cell population. (**h**) Electron dense particle similar in appearance to the cargo of NEB observed in connection with the ER. This event was observed in a cell exposed to oxidative stress. Scale bars: 2 μm (**a, c**), 1 μm (**b, d**), 200 nm (**e-h**). Abbreviations: N, nucleus; NEB, nuclear envelope budding; HMC-1, human mast cell line 1; black arrows indicate NEB events.

#### Supplementary figure 2 (Related to Figure 1)

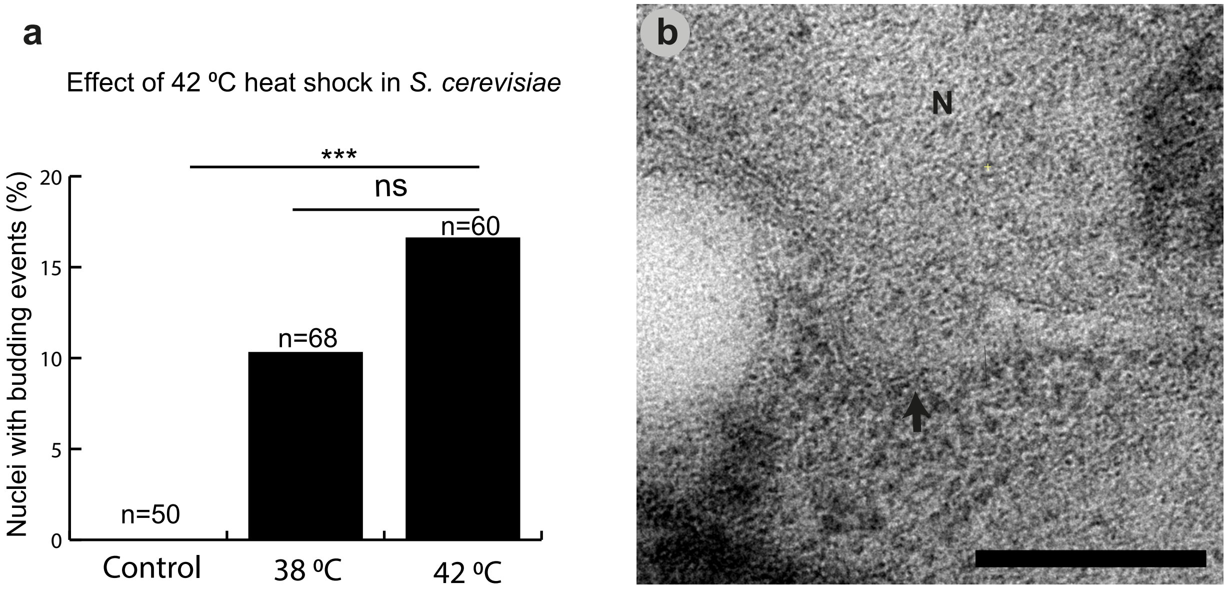


**Supplementary figure 2. Effect of 42⁰C heat shock in *S. cerevisiae*.** Cells were subjected to 42⁰C heat shock for 30 minutes to evaluate the influence of a higher temperature in the frequency of NEB. (**a**) The 42⁰C heat shock increased the frequency of NEB to a higher extend compared with control cells. (**b**) Representative micrograph of a NEB event found under the 42⁰C heat shock treatment. ***$P$<.001, ns no significant differences between groups. Scale bar: 200nm.

#### Supplementary figure 3 (Related to Figure 2)

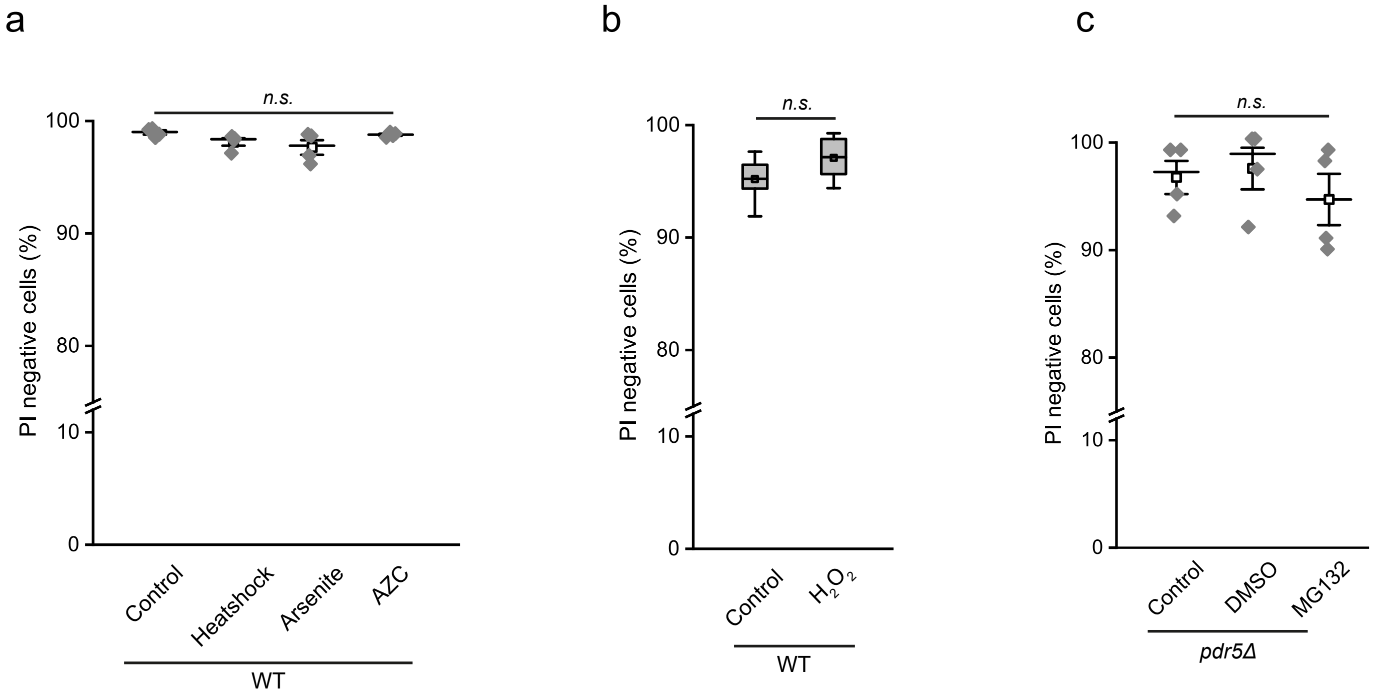


**Supplementary figure 3. Viability assay of cellular stressors.** For evaluating the condition of the cells after subjected to variant stressors, propidium iodide (PI) staining was performed. (**a-c**) Percentage of PI negative cells for each stressor and reagent that may have affected the viability of the cells. There was no significant difference between the control and the treated groups as illustrated in the graphs.

### Supplementary figure 4 (Related to Figure 3)

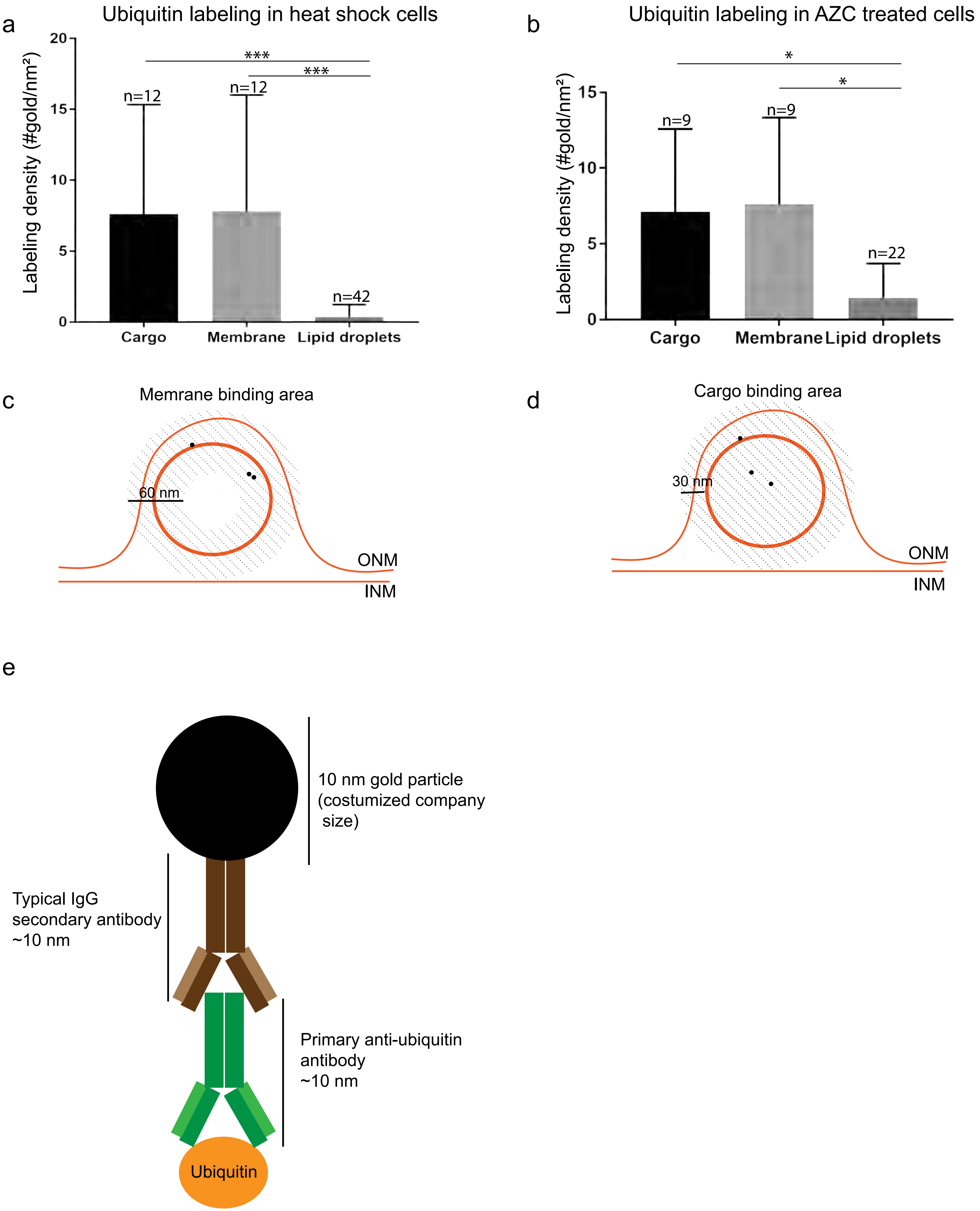

**Supplementary figure 4. Localization of immuno-labeling gold particles.** The localization of the gold particles for the ubiquitin assay, was characterized as ‘membrane binding’ or ‘cargo binding’ in accordance with their distance from the respective structures. (**a, b**) Labeling density of the ‘cargo binding area’, the ‘membrane binding area’ and lipid droplets (negative control). (**c, d**) Definition of the ‘cargo binding area’ and the ‘membrane binding area’. As the antibody sandwich that was used for this assay has a length of approximately 30 nm, the total area of the cargo including 30 nm of the surrounding area, was considered as the ‘cargo binding area’. In a similar manner, the area stretching 30 nm inwards and outwards of the cargo’s membrane was considered as the ‘membrane binding area’. (**e**) A graphical representation of the antibody sandwich and its approximate length. All graphs were generated using GraphPad Prism (version 8.1.2 (332)) software with the error bars representing the standard deviation. *$P$<.05, ****$P$<.0001 vs. lipid droplets (negative control).

#### Supplementary figure 5 (Related to Figure 6)

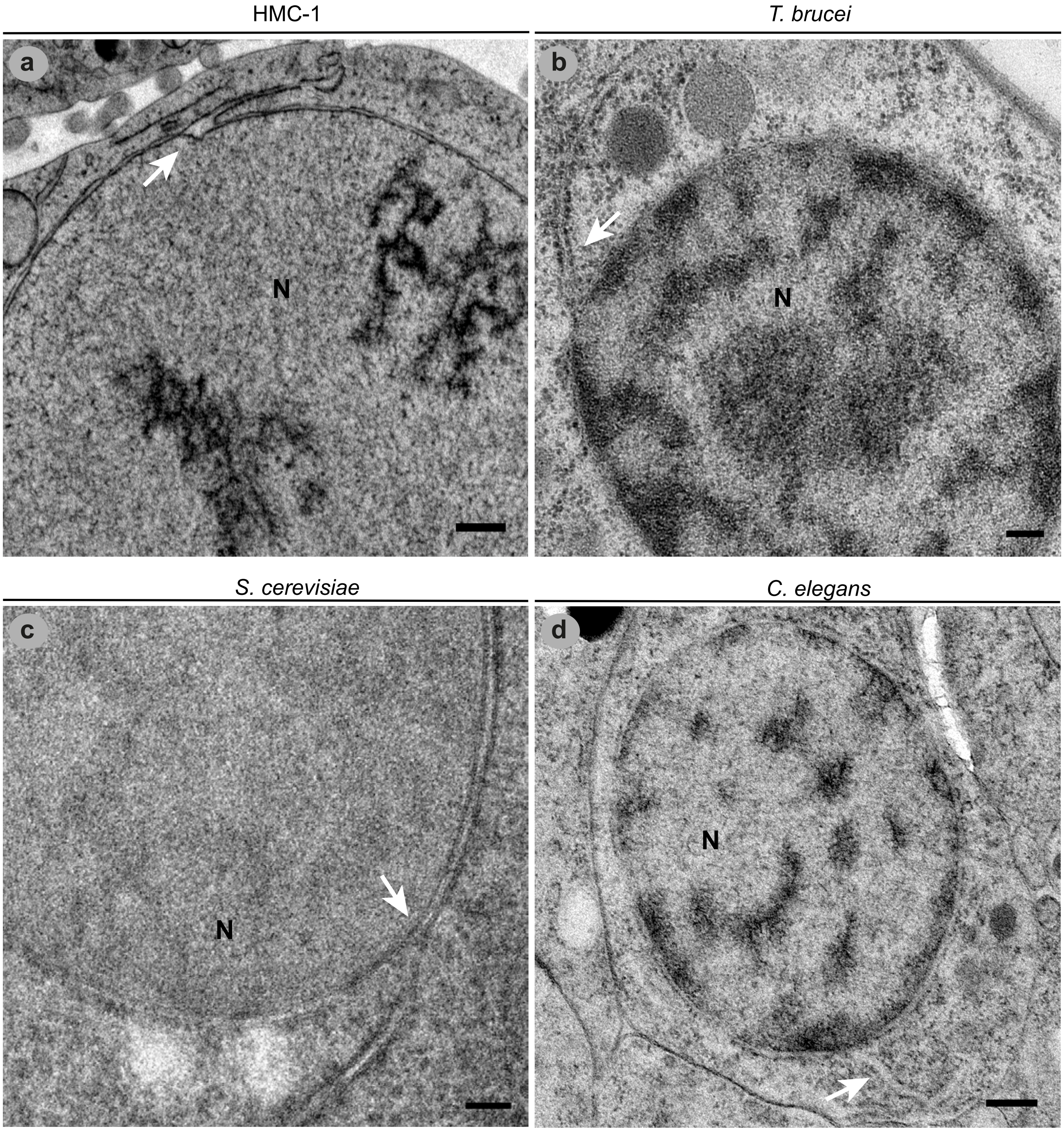


**Supplementary figure 5. Examples of NE-ER connections.** (**a-d**) NE-ER connections as they appear in 2D electron microscopy pictures in four different organisms. Their morphology is profoundly different from the one of the NEB events described in this study. Scale bars: 2 µm (**a**), 300 nm (**b**), 200 nm (**c**), and 500 nm (**d**). Abbreviations: NE, nuclear envelope; ER, endoplasmic reticulum; NEB, nuclear envelope budding; N, nucleus; white arrows indicate the NE-ER connection points.

**Supplementary figure 6 (Related to Figure 5)**

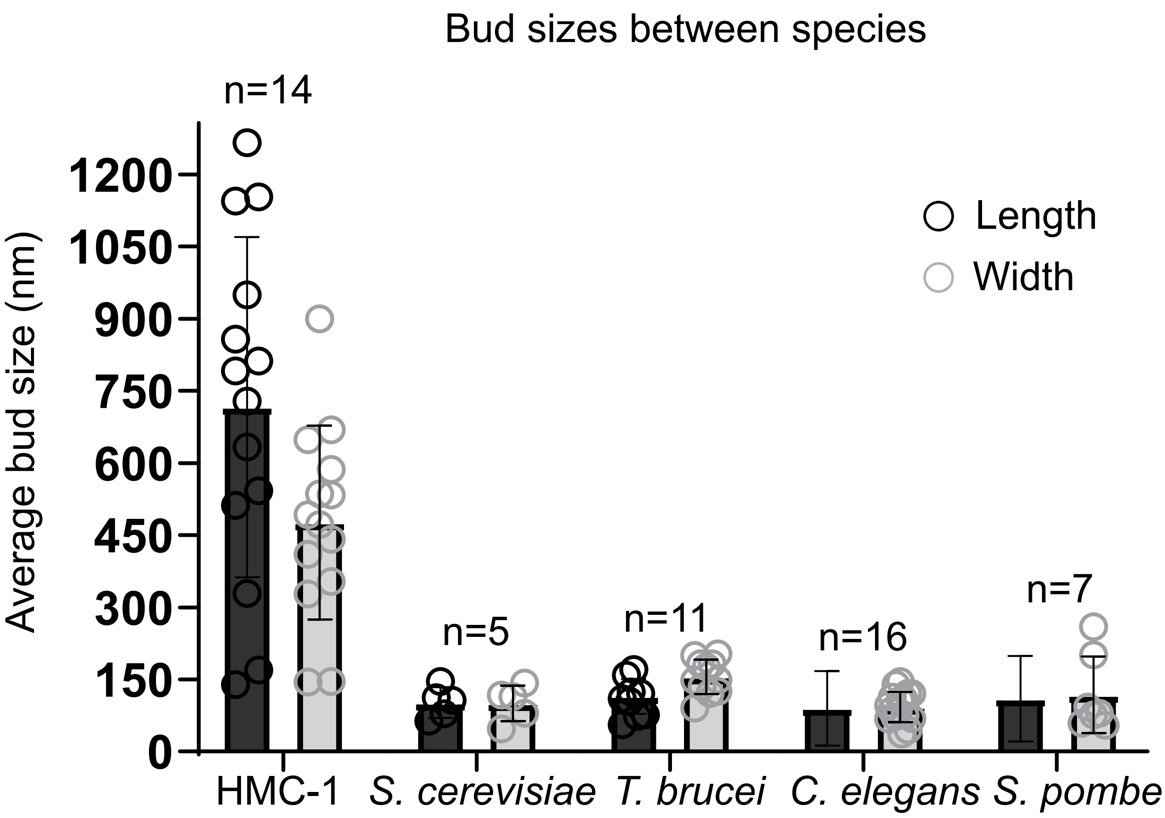


**Supplementary figure 6. Different sizes of NEB events.** Average sizes of nuclear buds (length and width) for each species. The n is equal to the total number of buds for the respective organism. Abbreviations: NEB, Nuclear envelope budding; HMC-1, Human mast cell line 1. The graph was generated using the GraphPad Prism (version 8.1.2 (332)) software with the error bars representing the standard deviation.

#### Supplementary figure 7 (Related to Figure 3)

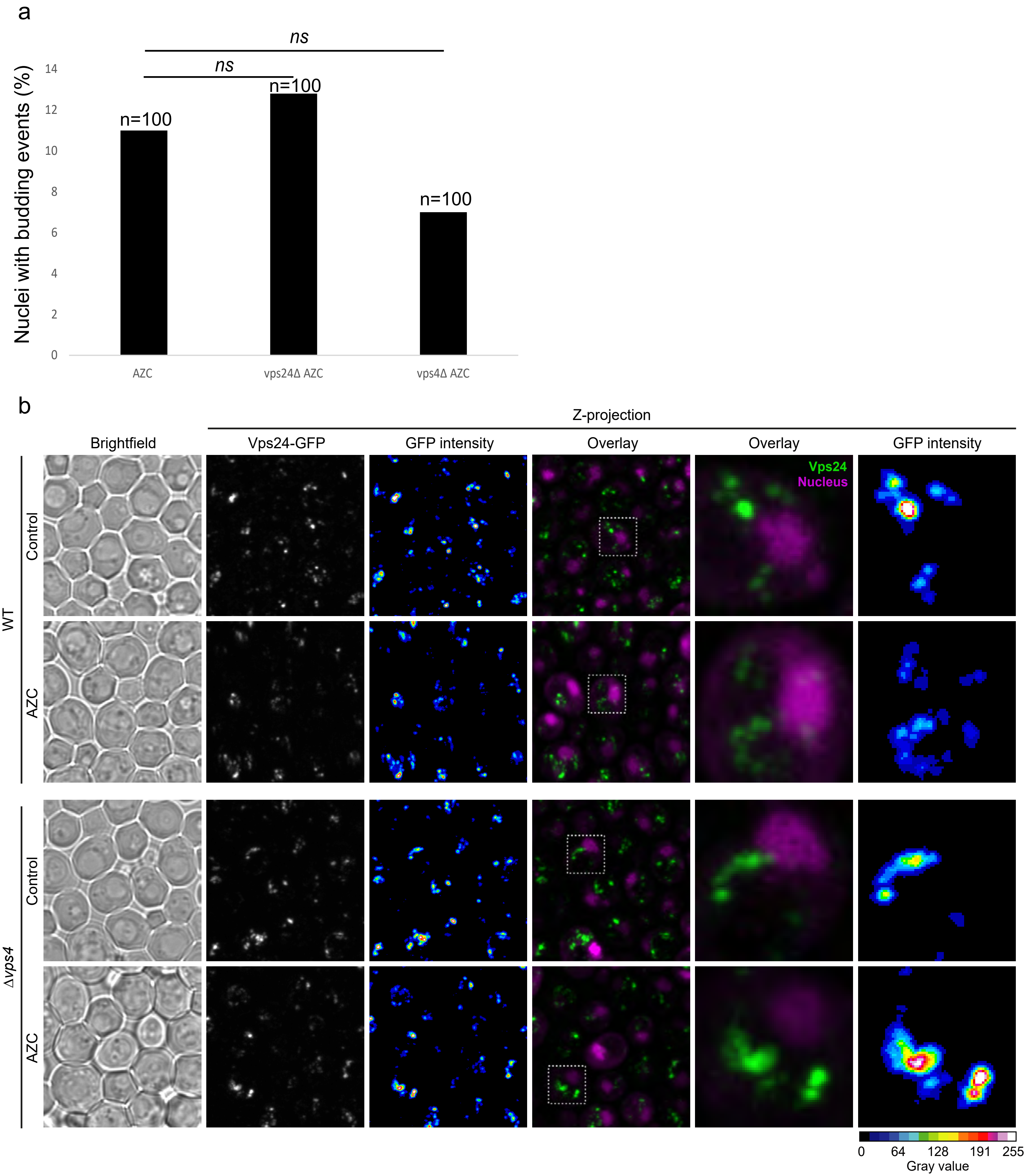

**Supplementary figure 7. No evidence for involvement of the ESCRT pathway in NEB.** (**a**) The possible involvement of the ESCRT pathway in the formation of NEB events, was examined through deletions of VPS24 and VPS4 genes. NEB frequency in the two deletion strains, *vps24Δ* and *vps4Δ* showed no significant difference to the control. In this experiment, all samples were treated with AZC for 90 minutes to enable the detection of a potential decrease in NEB frequency. (**b**) Confocal microscopy images of the Vps24-GFP protein both in wild type and *vps4Δ* strains show that the ESCRT protein labelled did not colocalize with the nucleus counterstained with DRAQ5. These experiments were performed both in normal and AZC treated conditions. Scale bars: 2μm.

### Supplementary Table 1

**Supplementary table 1.** List of organisms where NEB events have been previously observed, presented in a chronological order. The different terminologies used to describe the events are also included. *In this organism, the morphology of the budding events was somewhat different from the rest.

| Organism | Cell type | Year of publication | Terminology | Reference |
|---|---|---|---|---|
| *Drosophila melanogaster* | Salivary gland cells | 1955 | Membrane outpocketings | (Gay, 1955) |
| *Oryctolagus cuniculus* | Blastocysts | 1962 | Nuclear extrusion | (Hadek et al., 1962) |
| **Noctiluca scintillans* | unicellular | 1963 | Annulated vesicles | (Afzelius, 1963) |
| *Cricetulus griseus* | Embryo cell line $A_1$ | 1965 | Nuclear budding | (Longwell et al., 1965) |
| *Rattus norvegicus* | Fertilized oocytes | 1965 | Nucleolar extrusion | (D. Szollosi, 1965) |
| *Bos taurus* | Fibroblastic cells | 1965 | Nuclear buds | (Elston et al., 1965) |
| *Tradescantia bracteata* | Developing microspores | 1969 | Membrane bounded bodies | (Mepham et al., 1970) |
| *Podocarpus macrophyllus* | Haploid microspore tetrads | 1969 | Invaginations of nuclear envelope | (Aldrich et al., 1970) |
| *Lycopodium annotinum L.* | Immature spores | 1970 | Extrusions/bulges | (Gullvåg, 1970) |
| *Pinus banksiana* | Post meiotic microspores | 1970 | Folding of nuclear envelope | (Dickinson et al., 1970) |
| *Lilium longiflorum* | Young microspores | 1971 | Membrane-bound inclusions | (Dickinson, 1971) |
| *Drosophila melanogaster* | -Salivary gland cells<br>-midgut cells | 1987 | Pod-like infoldings of the nuclear envelope | (Hochstrasser et al., 1987) |
| *Mus musculus* | -Zygotes<br>-early embryos<br>-hybrid cells | 1988 | ‘Blebbing’ of nuclear envelope | (M. S. Szollosi et al., 1988) |
| *Drosophila melanogaster* | Larval muscle cells | 2012 | Nuclear envelope budding | (Speese et al., 2012) |
| *Strongylocentrotus purpuratus* | Embryonic cells | 2018 | Nuclear egress | (LaMassa et al., 2018) |
| *Saccharomyces cerevisiae* | Aging mitotic cells/NPC assembly mutants | 2019 | Herniations at the nuclear envelope | (Rempel et al., 2019) |
| *Drosophila melanogaster* | Salivary gland cells | 2020 | Nuclear envelope budding | (Verboon et al., 2020) |

| **Supplementary Table S2:** Summary of experimental observations | | | | | | | |
|---|---|---|---|---|---|---|---|
| | Measurements (N) | Observations (x) | x/N (%) | *p*-values are calculated against indicated control data [‡]Wilcoxon | [$]Fisher | [£]chi-squared | [€]t-test |
| **Figure 1a:** Effect of heat shock in *S. cerevisiae* | | | | | | | |
| **EDC** | | | | | | | |
| Control wild type cells | 76 | 27 | 35.5% | | | | |
| 5 min | 63 | 19 | 30.2% | 0.5064 | 0.8023 | 0.5032 | 0.5067 |
| 15 min | 72 | 60 | 83.3% | $< 10^{-4}$ | $< 10^{-4}$ | $< 10^{-4}$ | $< 10^{-4}$ |
| 30 min | 68 | 55 | 80.9% | $< 10^{-4}$ | $< 10^{-4}$ | $< 10^{-4}$ | $< 10^{-4}$ |
| 45 min | 70 | 57 | 81.4% | $< 10^{-4}$ | $< 10^{-4}$ | $< 10^{-4}$ | $< 10^{-4}$ |
| 90 min | 81 | 49 | 60.5% | 0.0018 | 0.0014 | 0.0018 | 0.0016 |
| **NEB** | | | | | | | |
| Control wild type cells | 337 | 8 | 2.4% | | | | |
| 5 min | 63 | 3 | 4.8% | 0.289 | 0.2426 | 0.2874 | 0.2886 |
| 15 min | 72 | 4 | 5.6% | 0.1475 | 0.1429 | 0.1465 | 0.1472 |
| 30 min | 68 | 7 | 10.3% | 0.0016 | 0.0059 | 0.0016 | 0.0016 |
| 45 min | 70 | 4 | 5.7% | 0.1337 | 0.1337 | 0.1327 | 0.1334 |
| 90 min | 81 | 4 | 4.9% | 0.2158 | 0.1864 | 0.2146 | 0.2156 |
| 15 min + 30 min + 45 min | 210 | 15 | 7.1% | 0.0069 | 0.0072 | 0.0069 | 0.0068 |
| **Figure 2d:** Effect of cell stress on NEB frequency | | | | | | | |
| **NEB events** | | | | | | | |
| Control wild type cells | 337 | 8 | 2.4% | | | | |
| Arsenite treated | 249 | 15 | 6.0% | 0.0247 | 0.0215 | 0.0245 | 0.0245 |
| Young cells | 204 | 14 | 6.9% | 0.0105 | 0.0107 | 0.0104 | 0.0104 |
| Old cells | 200 | 18 | 9.0% | 0.0006 | 0.0007 | 0.0005 | 0.0005 |
| Old cells vs. Young cells | | | | 0.4275 | 0.2708 | 0.4264 | 0.4277 |
| *$H_2O_2$ control | 197 | 3 | 1.5% | 0.5052 | 0.8369 | 0.5041 | 0.505 |
| *$H_2O_2$ treated | 204 | 12 | 5.9% | 0.0363 | 0.0332 | 0.0361 | 0.0361 |
| *$H_2O_2$ treated vs. $H_20_2$ control | | | | 0.0217 | 0.0189 | 0.0215 | 0.0214 |
| **Figure 2f:** Effect of AZC on NEB frequency | | | | | | | |
| **NEB events** | | | | | | | |
| Control wild type cells | 337 | 8 | 2.4% | | | | |
| AZC treated cells 30 min | 114 | 3 | 2.6% | 0.8788 | 0.5552 | 0.8775 | 0.8778 |
| [†]AZC treated cells 90 min | 100 | 22 | 22.0% | $< 10^{-4}$ | $< 10^{-4}$ | $< 10^{-4}$ | $< 10^{-4}$ |
| **Figure 3c:** Proteasome inhibition | | | | | | | |
| **NEB events** | | | | | | | |
| Control wild type cells | 337 | 8 | 2.4% | | | | |
| rpn4Δ | 100 | 15 | 15.0% | 0.0000 | 0.0000 | 0.0000 | 0.0000 |
| pdr5Δ + DMSO | 200 | 5 | 2.5% | 0.5689 | 0.9268 | 0.9277 | 0.9269 |
| pdr5Δ + MG132 | 200 | 14 | 7.0% | 0.0094 | 0.0089 | 0.009 | 0.0089 |
| pdr5Δ + MG132 vs. pdr5Δ + DMSO | | | | 0.0287 | 0.0344 | 0.0347 | 0.0344 |
| **Figure 4e:** Localization of NPC proteins | | | | | | | |
| **Immuno-gold labeling events** | | | | | | | |
| NPC unstressed cells | 100 | 81 | 81.0% | | | | |
| NPC aged cells | 95 | 63 | 66.3% | 0.0148 | 0.0197 | 0.0201 | 0.0196 |
| LDs unstressed cells | 83 | 15 | 18.1% | | | | |
| LDs aged cells | 100 | 8 | 8.0% | 0.0320 | 0.0371 | 0.0378 | 0.0372 |
| LDs vs NPCs unstressed cells | | | | $< 10^{-4}$ | $< 10^{-4}$ | $< 10^{-4}$ | $< 10^{-4}$ |
| LDs vs NPCs aged | | | | $< 10^{-4}$ | $< 10^{-4}$ | $< 10^{-4}$ | $< 10^{-4}$ |
| NEBs unstressed cells | 6 | 1 | 16.7% | | | | |
| NEBs aged cells | 10 | 1 | 10.0% | 0.6250 | 0.6963 | 1.0000 | 0.7192 |
| NEBs vs. NPCs unstressed cells | | | | 0.0238 | 0.0144 | 0.0152 | 0.0141 |
| NEBs vs NPCs aged cells | | | | $< 10^{-4}$ | $< 10^{-4}$ | $< 10^{-4}$ | $< 10^{-4}$ |
| **Supplementary Figure 1:** Effect of 42 $^{o}$C heat shock in S. cerevisiae | | | | | | | |
| **NEB events** | | | | | | | |
| Control wild type cells | 337 | 8 | 2.4% | | | | |
| Heatshock 38 °C | 68 | 7 | 10.3% | 0.0016 | 0.0059 | 0.0016 | 0.0016 |
| Heatshock 42 °C | 60 | 10 | 16.7% | 0.0000 | 0.0001 | 0.0000 | 0.0000 |
| Heat shock 42 °C vs. 38 °C | | | | 0.2891 | 0.212 | 0.2928 | 0.2928 |
| **Supplementary Figure 4:** Involvement of the ESCRT pathway in NEB | | | | | | | |
| **NEB events** | | | | | | | |
| *AZC treated wild-type cells 90 min | 100 | 11 | 11.0% | | | | |
| AZC treated Vps24Δ cells 90 min | 100 | 13 | 13.0% | 0.6658 | 0.7427 | 0.6634 | 0.6653 |
| AZC treated Vps4Δ cells 90 min | 100 | 7 | 7.0% | 0.3254 | 0.2297 | 0.323 | 0.3254 |

[†]Experiments involving $H_2O_2$ treated cells grew on different growth media and therefore an independent control was required.
*These experimental data were not merged to avoid potential batch-by-batch variations in AZC
[‡]https://se.mathworks.com/help/stats/ranksum.html
[$]https://se.mathworks.com/help/stats/fishertest.html
[£]https://se.mathworks.com/matlabcentral/fileexchange/45966-compare-two-proportions-chi-square
[€]https://se.mathworks.com/help/stats/ttest2.html

### Supplementary table 3

**Supplementary table 3.** Detailed list and information of all antibodies and reagents used in this study.

| | | |
|---|---|---|
| Antibodies | | |
| Anti-Nuclear Pore Complex Proteins antibody [Mab414] | Abcam | ab24609 |
| Anti-Ubiquitin antibody (for immuno-EM) | Abcam | ab19247 |
| | Abcam | ab6556 |
| Anti-GFP antibody | Abcam | ab69549 |
| Anti-Hsp104 antibody | | |
| Rabbit anti-mouse immunoglobulins | Agilent/Dako | E0433 |
| Gold-conjugated protein A | CMC UMC Utrecht | E1808 |
| Goat-anti-Rabbit IgG (H&L), 10nm | Electron Microscopy Sciences | Cat#25108 |
| Anti-Ubiquitin antibody (for WB) | Santa cruz | Sc-8017 |
| Chemicals | | |
| Uranyl acetate | SPI-CHEM | Lot#1221013 |
| HM20 non-polar Lowicryl | Polysciences Europe GmbH | Cat#15924-1 |
| Lead Nitrate | Merck | Cat#109969 |
| Sodium Citrate | Merck | Cat#111037 |
| Cationic gold particles (15 nm) | British Bio Cell | SKU#: EM.GC15 |
| Osmium tetroxide | TAAB | Cat#O014 |
| Fish skin gelatin | Sigma-Aldrich | Cat# 9000-70-8 |
| Sulfo-NHS-LC biotin | Thermo Fisher scientific | Cat#21335 |
| Streptavidin magnetic beads | Thermo Fisher scientific | Cat#21344 |
| Glutaraldehyde | Sigma-Aldrich Sweden AB | Cat#3802 |
| Sodium arsenite | Sigma-Aldrich | Cat# S7400 |
| 1-Hexadecene | Merck | Cat#629-73-2 |
| L-Azetidine-2-carboxylic acid (AZC) | Bachem | Cat# F-1281 |
| Hydrogen peroxide | Sigma-Aldrich | Cat# 7722-84-1 |
| Propidium iodide | Sigma-Aldrich | Cat#81845 |
| DRAQ5 | Abcam | Ab108410 |
| MG132 | Sigma-Aldrich | Cat#C2211 |

### Supplementary table 4

**Supplementary table 4.** Detailed list and information of all yeast strains and primers used in this study.

**Yeast strains**

| Strain | Genotype | Source |
|---|---|---|
| BY4741 | MATa, *his3*$\Delta$1, *leu2*$\Delta$0, *met15*$\Delta$0, *ura3*$\Delta$0 | Euroscarf |
| BY4741 *vps4*$\Delta$ | BY4741 *vps4*$\Delta$::hphNT1 | This study |
| BY4741 *vps4*$\Delta$ Vps24$^{GFP}$ | BY4741 *vps4*$\Delta$::hphNT1 *VPS24*-yeGFP-kanMX | This study |
| BY4741 Vps24$^{GFP}$ | BY4741 *VPS24*-yeGFP-kanMX | This study |
| BY4741 *vps24*$\Delta$ | BY4741 *vps4*$\Delta$::hphNT1 | This study |
| BY4741 *pdr5*$\Delta$ | BY4741 pdr5$\Delta$::hphNT1 | This study |

**Oligonucleotides**

| Modification | Oligonucleotides | PCR template |
|---|---|---|
| Deletion of *VPS4* | 5'-<br>ATGGAAGACAAAAATAAAGCAGCATAGAGTGCCTATAGTAGA<br>TGGG<br>GTACAAATGCGTACGCTGCAGGTCGAC -3'<br>5'-<br>TTTTTTATTTTCATGTACACAAGAAATCTACATTAGCACGTTAA<br>TCAA<br>TTGACTAATCGATGAATTCGAGCTCG-3' | pFA6a-hphNT1 |
| Control PCR<br>*VPS4* deletion | 5'-GTCGACCTGCAGCGTACG-3<br>5'- GATTCACATGTCGCCACTCCAGTC-3' | |
| C-terminal<br>tagging of *VPS24* | 5'-<br>CATTATTTATTCACTTATTTATTTATTTTCTTTGTACAGTCACAG<br>TAA<br>CACTCTAATCGATGAATTCGAGCTCG-3'<br>5'-GGAAGCAGATAGGATGGTAAATGAAATGCGTGAAAG<br>GCTGAGAGCTTTGCAAAACCGTACGCTGCAGGTCGAC-3' | pYM12 |
| Control PCR<br>*VPS24* tagging | 5'-CGAGCTCGAATTCATCGAT-3'<br>5'-GTGGGATCCAAAGACTGGAAC-3' | |
| Deletion of<br>*VPS24* | 5'-<br>ACCTTTAGTAGTTTGGGGGGCAGTTTTCTGGGCAATACAAAG<br>TTTA<br>CTTTTGATGCGTACGCTGCAGGTCGAC -3'<br>5'-<br>CATTATTTATTCACTTATTTATTTATTTTCTTTGTACAGTCACAG | pFA6a-hphNT1 |

### Supplementary table 5

**Supplementary table 5.** Summarized protocols on thin sections and tomography and immuno-EM samples.

| **Species** | **High-pressure freezing** | **Freeze substitution** | **Embedded plastic** | **Imaging** |
|---|---|---|---|---|
| *S. pombe* | Leica EM PACT1 | Long protocol (UA, GA, $OsO_4$) | HM20 | FEI Tecnai TF20 |
| *T. brucei* | Leica EM PACT2 | Short protocol (UA) | HM20 | Tecnai T12/Tecnai TF30 300 kV IVEM |
| *S. cerevisiae* | Wohlwend Compact 3 | Short protocol (UA) | HM20 | Tecnai T12/Tecnai TF30 300 kV IVEM |
| *C. elegans* | Wohlwend Compact 3 | Modified short protocol (UA) | HM20 | Tecnai T12 |
| HMC-1 | Leica EM PACT1 | Short protocol (UA) | K4M | Leo 912AB Omega TEM 120 kV |
| **Antibody** | **Dilution** | **Incubation time** | **Experiment** | **Order** |
| mAb414 | 1:50 | 2 hours | NPC labeling | First incubation |
| Rabbit anti-mouse immunoglobulins | 1:150 | 1 hour | NPC labeling | Second incubation |
| 10nm gold-conjugated protein A | 1:70 | 30 minutes | NPC labeling | Third incubation |
| Ab19247 | 1:20 | 2 hours | Ubiquitin labeling | First incubation |
| 10nm gold Goat-anti-Rabbit IgG | 1:20 | 1 hour | Ubiquitin, GFP, Hsp104 labeling | Second incubation |
| Ab6556 | 1:5, 1:10, 1:30 | Overnight, 4°C | GFP labeling | First incubation |
| Ab69549 | 1:100 | Overnight, 4°C | Hsp104 labeling | First incubation |
